# Site-specific hyperphosphorylation of tau inhibits its fibrillization *in vitro*, blocks its seeding capacity in cells, and disrupts its microtubule binding; Implications for the native state stabilization of tau

**DOI:** 10.1101/772046

**Authors:** Mahmood Haj-Yahya, Pushparathinam Gopinath, Kolla Rajasekhar, Hilda Mirbaha, Marc. I. Diamond, Hilal A. Lashuel

## Abstract

The consistent observation of aggregated phosopho-tau in the pathology of Alzheimer’s disease and other tauopathies has contributed to the emergence of a model where hyperphosphorylation of tau causes its disassociation from microtubules and subsequent pathological polymerization. However, the large number of possible phosphorylation sites in tau and lack of robust methods that enable the preparation of homogeneously phosphorylated tau species have made it difficult to validate this model. Herein, we applied a total chemical synthetic approach to site-specifically phosphorylate single (pS356) or multiple (pS356/pS262 and pS356/pS262/pS258) residues within the microtubule binding repeat domain (MTBD) of tau and show that hyperphosphorylation within the microtubule MTBD inhibits K18 tau 1) aggregation in vitro; 2) its seeding activity in cells, and 3) its ability to promote microtubule polymerization. The inhibition increased with the number of phosphorylated sites, with phosphorylation at S262 having the strongest effect. On the basis of these findings, we propose that targeting the kinases that regulate phosphorylation at these sites could provide a viable strategy to stabilize the native state of tau and inhibit its aggregation. Taken together, our results argue against the pathogenic hyperphosphorylation hypothesis and underscore the critical importance of revisiting the role of site-specific hyperphosphorylation of tau in regulating its function in health and disease.

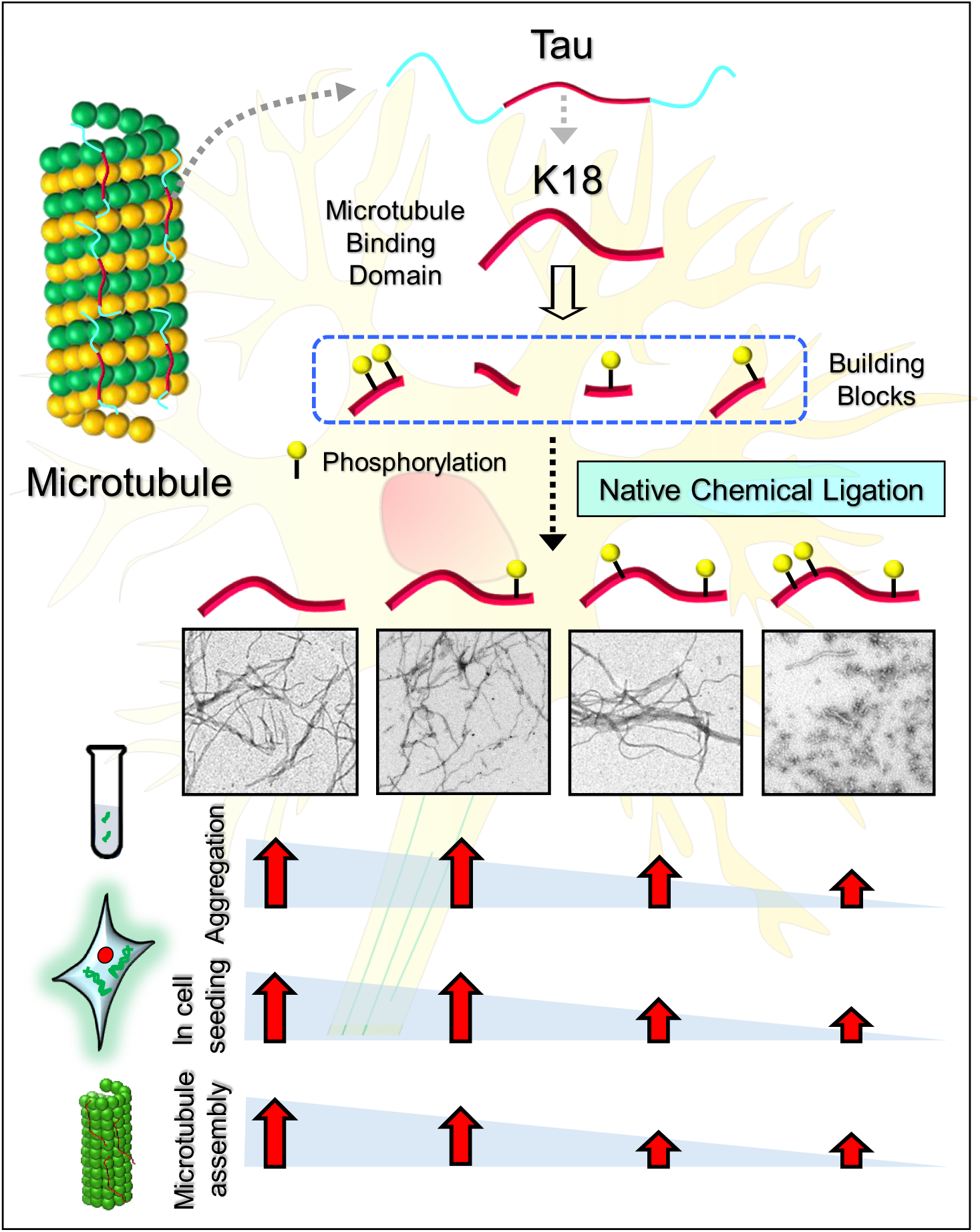
Table of content.

## Introduction

The microtubule binding protein tau is the primary component of the neurofibrillary tangles (NFTs) found in the brain of Alzheimer’s disease (AD) patients^1^. Aggregated tau is also present in other pathological protein aggregates that characterize several other neurodegenerative diseases, including progressive supranuclear palsy, corticobasal syndrome, frontotemporal dementias, and chronic traumatic encephalopathy (CTE), collectively known as tauopathies. Although initially thought to be a secondary and downstream effect of amyloid pathology in AD, increasing evidence from human clinical trials,^2, 3^ animal models^4, 5^ and longitudinal imaging studies^6, 7^ points towards a central and causative role for tau aggregation in the initiation and progression of AD^8^. The exact mechanisms by which tau contributes to neurodegeneration in tauopathies, the role of phosphorylation and other posttranslational modifications (PTMs) on tau aggregation, pathology spreading and toxicity, and the nature of the toxic species remain poorly understood.

Several aspects of tau function in health and disease are regulated by PTMs including phosphorylation, acetylation, ubiquitination and truncation^6^. However, deciphering the tau PTM code has proven to be challenging for the following reasons: 1) More than seven types of PTMs have been described in different regions of tau; 2) For each modification, several sites are reported to be modified. For example, there are at least 85 putative phosphorylation sites (80 serine/threonine and 5 tyrosine residues). Among those, 42 tau phosphorylation sites have been experimentally detected^6^. 3) With the exception of truncations, serine/threonine phosphorylation and acetylation, no other PTMs could be partially mimicked by introducing natural mutations, thus making it difficult to approximate the effect of such modifications on the regulation of the structure, function and/or aggregation propensity of tau; and 4) The existing tools and methods allow only for the specific investigation or manipulation of one PTM at a time while tau at any given time is subjected to multiple PTMs. Towards addressing these challenges, we recently developed a semisynthetic strategy that enables the site-specific introduction of single or multiple PTMs within residues 246-441 of tau and used this approach to elucidate the effect of acetylation at K280 on tau aggregation and tau-mediated tubulin polymerization^8, 9^. In this work, we have leveraged these advances to optimize a synthetic method for producing the K18 fragment, consisting of the microtubule binding repeat domain (MTBD) of tau. We applied this method to investigate the effects of phosphorylation at four residues of the MTBD on microtubule binding, *in vitro* aggregation, and seeding in cell models.

We chose the K18 fragment to investigate the role of tau phosphorylation for the following reasons. 1) It contains all four microtubule binding repeats (R1, R2, R3 and R4) that are involved in the binding of tau to microtubules and bear several of the disease-causing mutations associated with tauopathies^10^; 2) It contains several sequence motifs and PTM sites (Fig. 1) that have been shown to strongly influence tau aggregation and pathology formation, tau tubulin binding and microtubule assembly, stability and dynamics, although other regions in the protein also influence the dynamics of these processes (Fig. 1); 3) Cryo-EM studies of AD-derived PHFs^11^ and CTE-derived tau filaments^12^ have revealed that the R3 and R4 repeats constitute the core of the PHFs in the AD brain, similarly cryo-EM structure derived from narrow pick filaments (NPFs) of Pick disease brain revealed that the R1, R3 and R4 repeats constitute the core of the NPFs ^13^; and 4) The K18 fragments reproduce many of the key features of tau aggregation, exhibit rapid aggregation *in vitro*^14^, and seed more efficiently in cells^15^ compared to the full-length tau protein^16^. Therefore, the development of a synthetic strategy that enables site-specific modifications in the K18 fragment is highly relevant for elucidating the sequence and structural determinants of tau structure and aggregation.

**Fig. 1.**
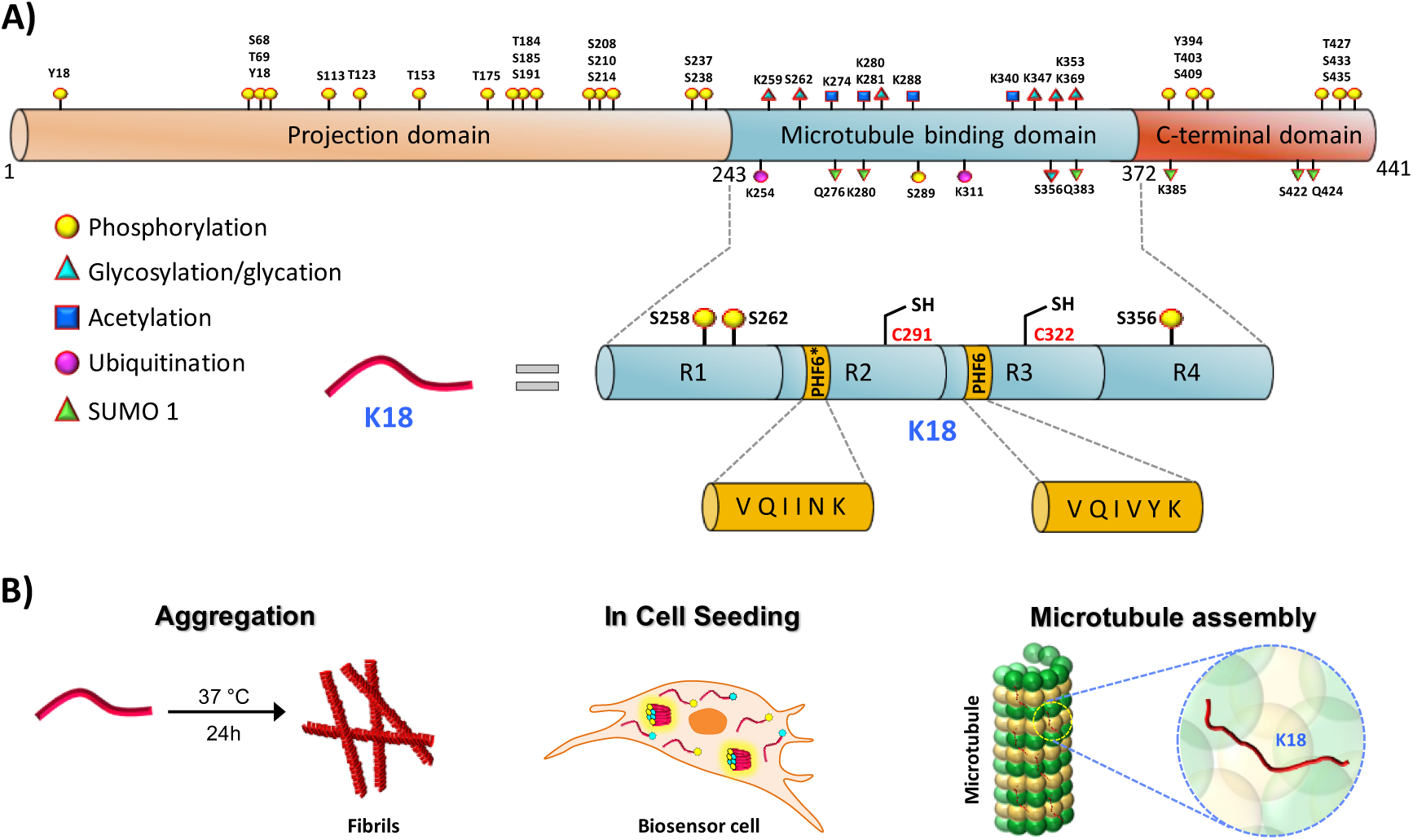
(A) Schematic representation of the longest tau isoform (2N4R), which contains an N-terminal projection domain, microtubule binding domain, and a C-terminal domain (PTMs highlighted). The microtubule binding domain is composed of four microtubule binding repeats, R1, R2, R3, and R4. Two hexapeptides, PHF6* and PHF6 in the R2 and R3 repeats, play critical roles in tau fibrillization. Among the 85 putative phosphorylation sites (80 serine/threonine and 5 tyrosine residues) on tau, 28 sites were identified to be exclusively phosphorylated in AD brains. In the case of MTBD, the sites S258, S262, S289 and S356, which occur in the MTBD, are heavily phosphorylated in AD pathology. (B) Schematic depiction illustrating the different experimental approaches used in this study to elucidate the effect of single and multiple phosphorylation states on the functional (tubulin assembly) and pathological (aggregation and seeding) aspects of K18.

Phosphorylation is one of the most actively investigated tau PTMs^6^ because pathological tau aggregates observed in the AD brain are heavily phosphorylated^17^. Furthermore, phosphorylation at several specific sites^18^ has correlated with the progression of tau aggregation and are thus used as markers of pathology. These observations have led to the hypothesis that phosphorylation disrupts tau microtubule interactions and enhances its misfolding and aggregation. Previous attempts to test this hypothesis have relied on the use of phosphomimetic mutations^19, 20^, which do not fully recapitulate all the aspects of phosphorylation, or kinases^17^ that phosphorylate tau at multiple positions with variable efficiency. The latter approach gives rise to mixtures of tau species with different phosphorylation patterns, rendering it impossible to elucidate the role of specific phosphorylation sites or patterns on the regulation of tau aggregation and microtubule binding.

K18 consists of 16 potential phosphorylation sites (12 serines/4 threonines); among them, 12 can be phosphorylated under physiological conditions^21, 22^. However, the remaining 4 sites (S256, S262, S289, and S356) are associated with AD. Several *in vitro* studies have shown that members of different kinase families phosphorylate K18 at multiple residues^23, 24^. Phosphorylation at S256, S262, and S356 residues is of key interest due to its occurrence in PHFs^25^. Kinases such as AMP-activated protein kinase, microtubule affinity-regulating kinase (MARK), calcium calmodulin kinase II and checkpoint kinase 1 have also been shown to phosphorylate the residues S256, S262, and S356^23, 26, 27^. However, these kinases lack site specificity and phosphorylate tau at several other residues.

Herein, we describe a novel, total chemical synthetic strategy that allows the preparation of site-specific and homogeneously modified forms of K18 (consisting of 130 amino acids (AAs)). We used this method to determine the effect of K18 phosphorylation at both single (pS356) and multiple residues (pS262, pS356, or pS262, pS258, pS356) on aggregation *in vitro*, tubulin assembly and seeding activity in a cellular model of tauopathy. Our studies demonstrate that phosphorylation within the MTBD inhibits rather than promotes tau fibrillization. We also show that the phosphorylation at S262 is a major contributor to disrupting tau-microtubule binding. On the basis of these findings and the modest effect of phosphorylation at S262 on K18 aggregation, we propose that inhibitors of kinases that regulate phosphorylation at this site could provide a viable strategy to stabilize the native state of tau and inhibit its aggregation.

## Results

### Design and total synthesis of site-specifically phosphorylated K18 proteins

Our strategy for the chemical synthesis of K18 is based on a three-fragment approach utilizing native chemical ligation (NCL)^28^ (Scheme 1). To maintain the native sequence of K18, we designed our strategy by taking advantage of the two native cysteine residues (Cys^291^ & Cys^322^), to serve as ligation sites. The three fragments were synthesized by Fmoc-solid phase peptide synthesis (Fmoc-SPPS). Since SPPS is limited to peptides that contain ≤ 40 AAs^28^, we first tested the feasibility of synthesizing the longest fragment tau **1** (322-372), which is composed of 51 AAs.

**Scheme 1.**
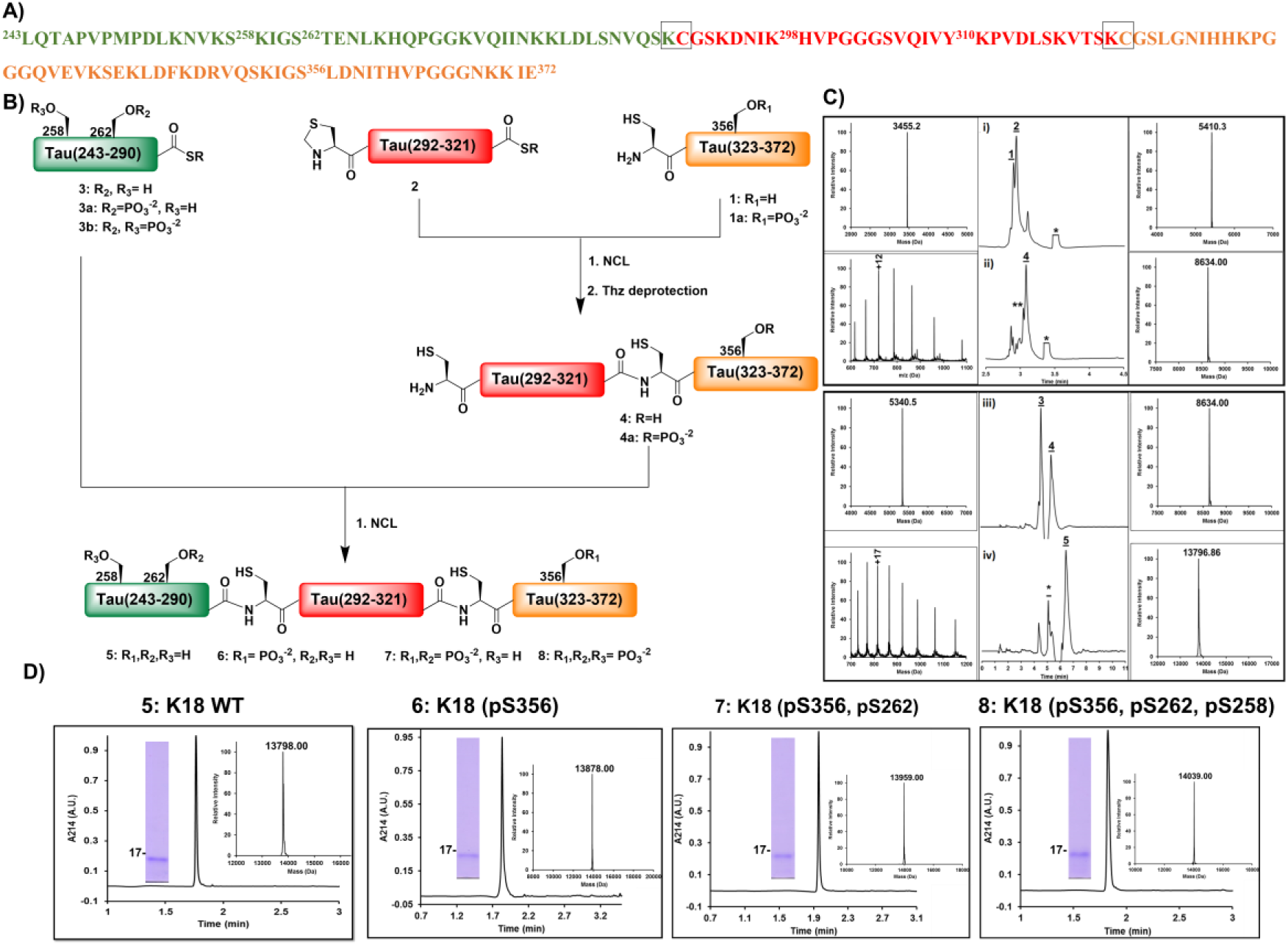
Strategy for the total chemical synthesis of the WT, and mono-, di-, and triphosphorylated K18. A) K18 amino acid sequence with the three fragments shown in green, red and orange and the ligation sites highlighted by gray boxes. B) Schematic representation of the synthesis of hyperphosphorylated K18 from the three synthetic fragments. C) Analytical RP-HPLC traces and mass spectrometry (ESI-MS) analysis of the ligation reactions between the different fragments of K18. Ligation of fragments 1 and 2 i) at 0 h and ii) at 4.5 h followed by in situ thiazolidine deprotection. Ligation of fragments 3 and 4 iii) at 0 h and iv) 2 h. D) Characterization of the purified K18 proteins by analytical RP-HPLC, ESI-MS, and SDS-PAGE.

During our initial efforts to prepare fragment **1** using rink amide resin (loading 0.55 mmol/g) and HCTU as the coupling reagent, we observed deletions of several amino acids, especially glycine (Gly) residues in regions 333-335 and 367-369, resulting in poor yields. To overcome these challenges, we used a low-loading rink amide resin (0.27 mmol/g), double coupling of all AAs, and incorporated pseudoproline dipeptides at Ile^360^-Thr^361^ and Lys^340^-Ser^341^. Under these conditions, we succeeded in obtaining fragment **1** in good yield (25%) and high purity (Fig. S1). Next, we synthesized fragments **2** (291-321) and **3** (244-290). For both fragments, C-terminal thioesterification was performed using the 3-(Fmoc-amino)-4-(methylamino) benzoic acid (Fmoc-MeDbz) linker as a precursor, as described previously by Dawson and coworkers^29^. After introducing the last amino acid coupled with N-terminal Boc protection, the C-terminus of the peptide was activated by acylation with p-nitrophenyl chloroformate and cyclized using a solution of 0.5 M DIEA in DMF (Figs. S3 and S4). As fragment **2** has both an N-terminal and C-terminal Cys thioester, the N-terminal Cys residue was introduced as thiazolidine (Thz) to avoid undesired inter- and/or intramolecular ligations.

With the three tau peptide fragments in hand, we then started the assembly of K18 via NCL of fragments **1** and **2**. The ligation reaction was complete after 4.5 h, and the resulting crude product bearing an N-terminal Thz was then treated with 0.2 M methoxyamine hydrochloride for 12 h at pH 4 to free the N-terminal cysteine required for the subsequent ligation. After completion of the Thz deprotection, the desired product was purified by RP-HPLC to provide fragment **4** (291-372) in ∼17% overall yield (Fig. S7). Next, the second ligation was carried out by dissolving thioester peptide **3** (243-290)-SR and fragment **4** (291-372) in 8 M urea followed by the addition of 50 mM tris(2-carboxyethyl)phosphine (TCEP) as a reducing agent and 50 mM 4-mercaptophenylacetic acid (MPAA) as a thiol additive. The purification of the final product proved to be difficult due to its hydrophilicity, which led to its coelution with MPAA. To circumvent this problem, we used the volatile alkyl thiol trifluoroethanethiol (TFET)^30^, which can be easily removed after completion of the ligation reaction by evaporation with nitrogen gas for 30 min. The final product, Wt K18 **5**, was purified by RP-HPLC in high purity and ∼20% yield (Fig. S8).

To verify that synthetic K18 adopts the correct conformation and is functional, we assessed and compared its secondary structure and tubulin polymerization activity to that of the recombinant K18 produced in E coli as described previously^31, 32^. Both the synthetic and recombinant K18 exhibited virtually identical CD spectra (Fig. 2B) and tubulin polymerization activity (Fig. 2C)^33^. Next, we achieved the synthesis of K18 phosphorylated at single (pS356), double (pS356, pS262) or multiple sites (pS356, pS262, pS258), following the same synthetic strategy (Figs. S9-S12) as was used for the WT K18, using one or a combination of the following phosphorylated peptide fragments: peptide **1a** (322-372, pS356), **3a** (243-290, pS262)-SR and **3b** (243-290, pS262, pS258)-SR (Figs. S2, S5 and S6).

**Fig. 2.**
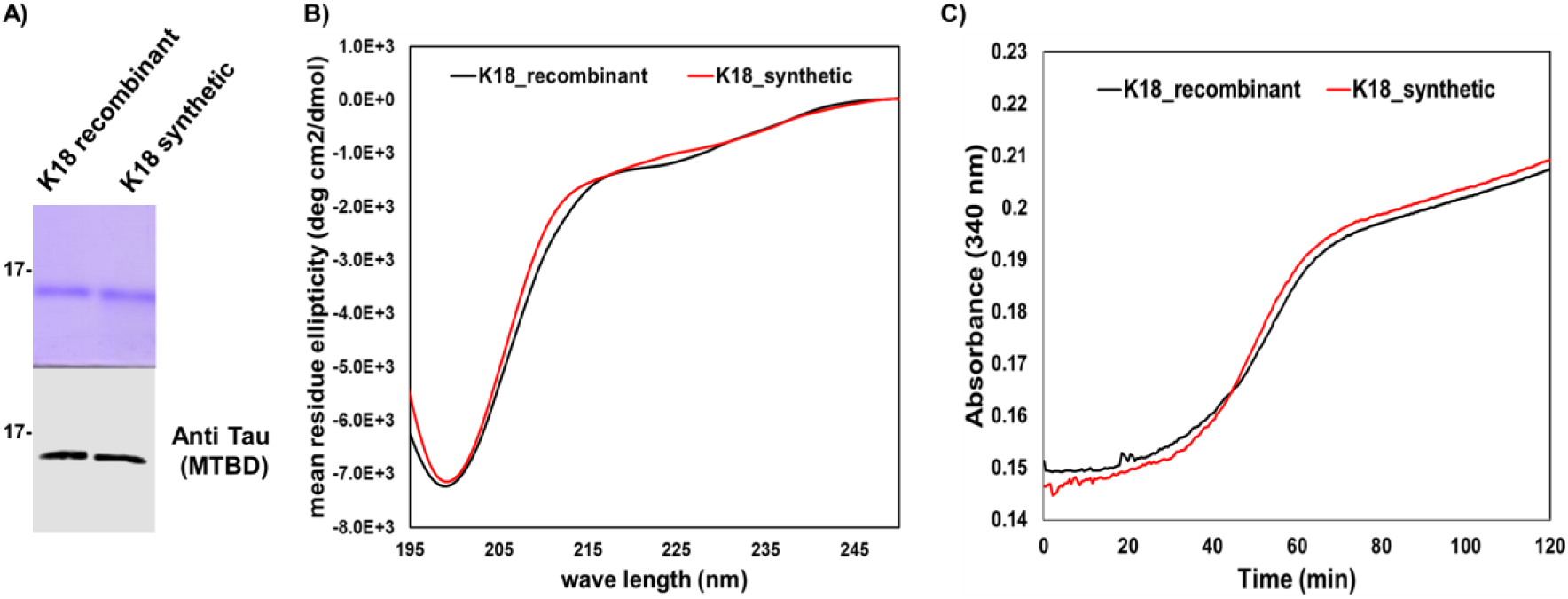
Characterization of synthetic and recombinant WT K18. A) SDS-PAGE analysis (upper panel) of WT recombinant and synthetic K18 as well as a western blot using the tau specific antibody (Polyclonal rabbit anti-total tau antibody, generated in house) (lower panel). B) Circular dichroism spectra of the WT recombinant (10 μM, black) and synthetic (10 μM, red) K18. C) Tubulin polymerization assay in the presence of synthetic and recombinant WT K18.

### An increase in phosphorylation decreases the aggregation of K18

To study the effect of site-specific phosphorylation on the aggregation of K18, synthetic K18 proteins (10 µM) bearing one (pS356), two (pS356, pS262) or three (pS356, pS262, pS258) phosphoserine residues were incubated at 37 °C in the presence of the polyanion heparin, which promotes fibril formation^32^. The kinetics of fibril formation were assessed by monitoring the changes in thioflavin S (ThS) fluorescence over time. As shown in Fig. 3A, the WT K18 protein reaches the aggregation plateau within 4-5 h. Phosphorylation of serine 356 (K18_pS356) did not significantly affect the aggregation kinetics of K18 but resulted in significantly lower ThS values compared to the unmodified K18. In contrast, phosphorylation of both S262 and S356 significantly delayed but did not inhibit the aggregation of K18, as observed by the longer nucleation phase and slower growth slope. Interestingly, phosphorylation at three residues (pS356, pS262 and pS258) completely abolished fibril formation, as no increase in the ThS fluorescence was observed over time, underscoring the inhibitory impact of phosphorylation on K18 aggregation.

**Fig. 3.**
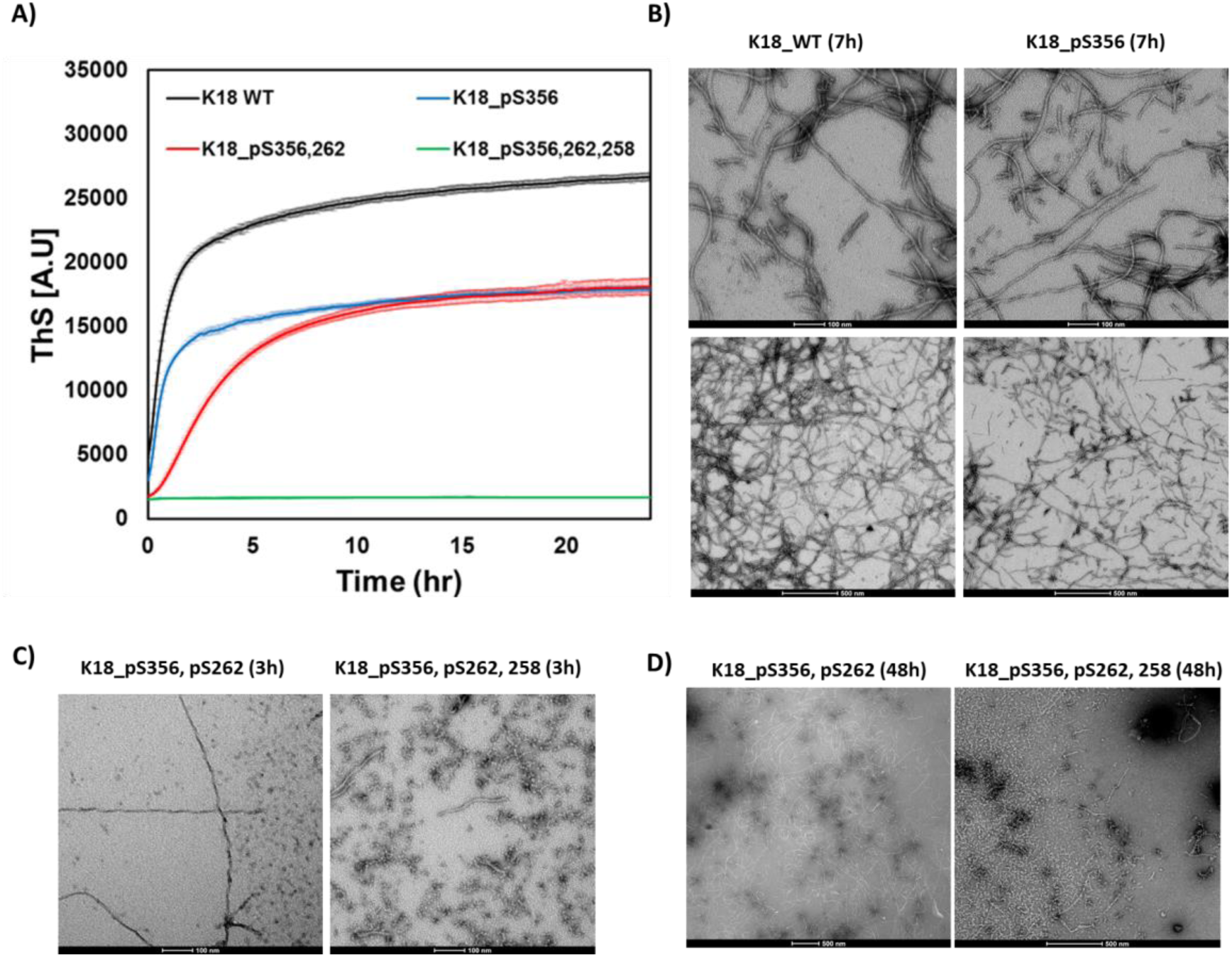
*In vitro* aggregation of WT and phosphorylated K18 proteins. A) Aggregation kinetics measured by ThS fluorescence at 490 nm (mean ± SEM, n=3). B) Schematic representation of WT, mono-, di- and triphosphorylated K18 constructs. C) TEM images of WT and K18_pS356 after 7 h (scale bars are 100 nm (upper panels) and 500 nm (lower panels)). D) TEM images of K18_ pS356, pS262 and K18_ pS356, pS262, pS258 after 3 h (scale bar is 100 nm) and E) 48 h of incubation (scale bar is 500 nm).

To further corroborate the ThS data and assess the effect of mono-, di- and triphosphorylation on K18 fibril morphology, the aggregates were examined by transmission electron microscopy (TEM). As shown in Fig. 3B-D, both mono- and diphosphorylated K18 rapidly form fibrillar aggregates similar to those previously reported for K18. In the case of the diphosphorylated K18, both fibrils and oligomers are observed at 3 h (Fig. 3C, left panel), while only fibrils are detected at longer incubation times (48 h, Fig. 3D left panel). The lower ThS values for the phosphorylated K18 proteins suggest that the phosphorylated fibrils might exhibit lower binding to ThS. Interestingly, triphosphorylated K18 also self-assembles into oligomeric structures within the first 3 h (Fig. 3C, right panel), but these aggregates persist over time and do not go on to form fibrils; however, we observed a very small amount of short fibrils at long incubation times (Fig. 3D, right panel). The negligible number of fibrils observed by EM likely explains the absence of a ThS signal in the kinetic aggregation assay (Fig. S14). These findings, derived from the homogeneously phosphorylated synthetic proteins, counter the conventional thinking that associates hyperphosphorylation with the induction or promotion of tau aggregation^7^.

### An increase in phosphorylation decreases the seeding efficiency of K18

To investigate the effect of phosphorylation at these sites on the seeding activity of K18, we used the previously described HEK293T biosensor cell reporter lines^34^. As depicted in Fig. 4A, these cells stably express the K18 4R tau repeat domain (RD, K18) containing the disease-associated mutation P301S fused to either cyan or yellow fluorescent protein (RD-CFP/RD-YFP, respectively). RD-CFP and RD-YFP exist exclusively as monomers unless the cells are exposed to exogenous tau aggregates, which are taken up micropinocytosis^35^. Here, we introduced the aggregates directly into the cells using a cationic lipid transfection reagent (Lipofectamine), which increases the assay sensitivity by approximately 100-fold^34^. The uptake of exogenous aggregates induces intracellular aggregation of RD-CFP/YFP, which creates a fluorescence resonance energy transfer (FRET) signal. Within a population of cells, the percentage that contain intracellular aggregates can be accurately measured using FRET-flow cytometry. To determine the titer of tau seeding activity, a FRET value is derived for a cell population using the integrated FRET density (IFD), which is the product of the percent of positive cells and the mean fluorescence intensity of the FRET-positive cells^34^.

**Fig. 4.**
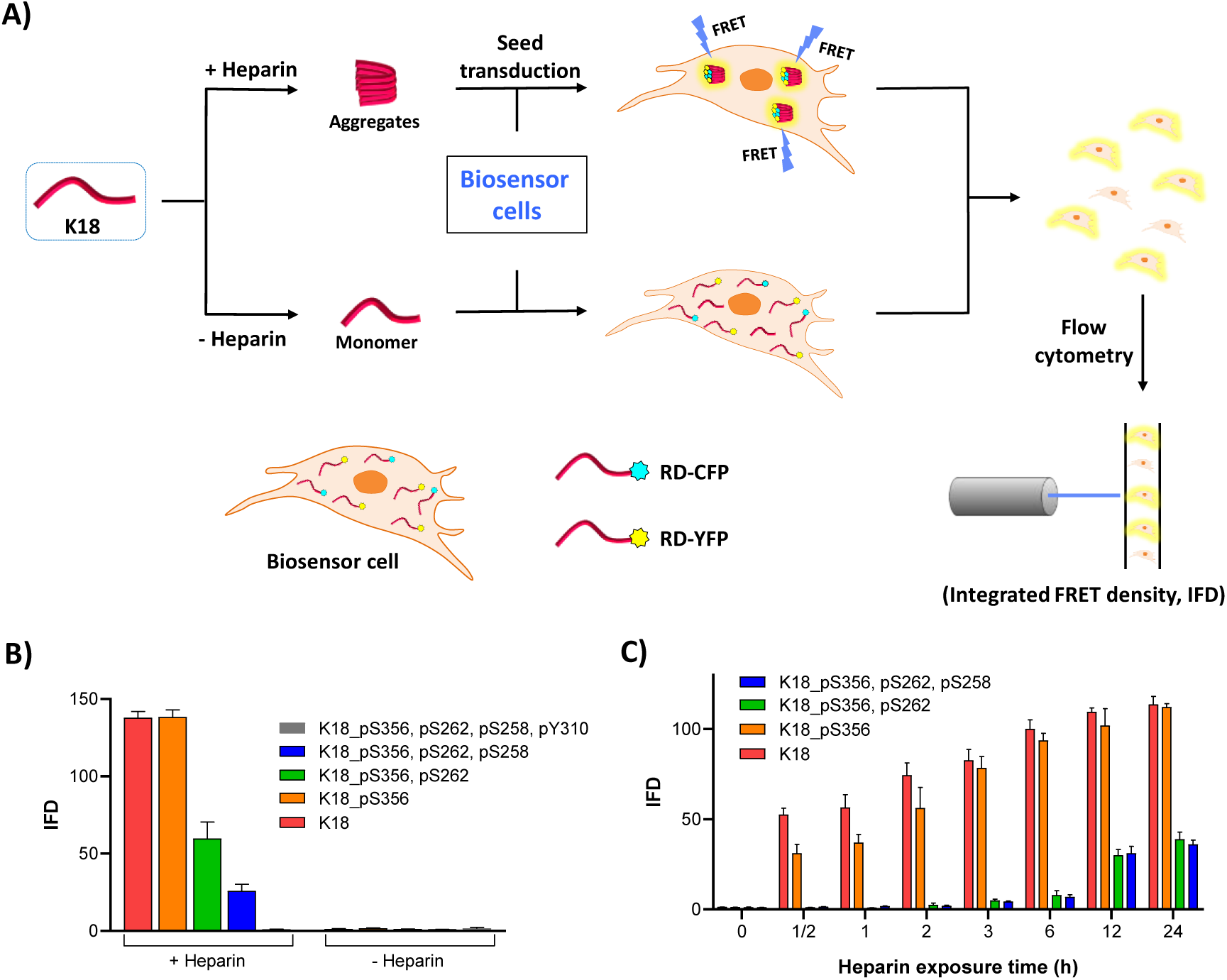
A) Schematic representation of the FRET-based seeding assay using HEK293T biosensor cell reporter lines. Heparin-induced aggregates are transfected into the biosensor cells using Lipofectamine. Aggregation seeded by the K18 proteins is detected as FRET emission and quantified using IFD. B) IFD measured following transduction of WT, mono-, di- tri and tetraphosphorylated K18 previously incubated with or without heparin for 24 h at 37 °C. C) Effect of different heparin exposure times with K18 proteins prior to transfection of the aggregates.

When the WT and phosphorylated K18 variants were incubated in the absence of heparin for 24 or 72 hours at 37°C or 25°C and then added to the biosensor cell reporter line, we did not observe seeding activity (Fig. 4B). This is consistent with the fact that tau does not spontaneously form fibrils in the absence of polyanionic inducers. However, when WT and K18_pS356 were incubated in the presence of heparin for 24 or 72 hours at 37°C, significant seeding activity was observed, while constructs bearing double (pS356, pS262) or triple (pS356, pS262, pS258) phosphorylated sites showed significantly lower seeding activity (Figs. 4B and S15). Interestingly, the tetraphosphorylated K18 protein bearing one additional phosphate group at Y310, which occurs within one of the key aggregation motifs (PHF6, Fig. 1), did not exhibit any seeding capacity (Fig. 4B).

When the fibrils were incubated in the presence of heparin for shorter periods before the addition to the biosensor cells, we observed that WT K18 and K18_pS356 have different seeding activity when added to the biosensor cells at shorter incubation times (0-12 h, Fig. 4C), whereas WT K18 developed seeding activity more rapidly than K18_pS356 (Fig. S16). Together, these findings suggest a direct correlation between the *in vitro* aggregation propensity of the K18 proteins and their seeding activity in the cell-based biosensor assay.

### Phosphorylation at S262 disrupts tau binding to microtubules

One of the key physiological functions of tau is its role in promoting tubulin polymerization and the stabilization of microtubules.^18^ Several studies have shown that phosphorylation at single or multiple residues in the MTBD could regulate or negatively influence tubulin polymerization. However, all of these studies were based on experimental approaches that did not allow for efficient and site-specific phosphorylation of tau or relied on the use of phosphomimetic mutations, which do not reproduce the effects of bona fide phosphorylation. To assess the effect of authentic phosphorylation at single or multiple residues within the MTBD on the modulation of microtubule assembly, we performed a tubulin polymerization assay^36^. The tubulin protein (40 µM) was incubated with K18 proteins (30 µM) bearing one (pS356), two (pS356, pS262) or three (pS356, pS262, pS258) phosphoserine residues at 37 °C in the presence of GTP, as previously described^37, 38^. The kinetics of the MT assembly were assessed by monitoring the changes in absorbance at 350 nm over time. As expected, the native K18 protein accelerated the rate of microtubule polymerization (Fig. 5A)^36^. In contrast, the phosphorylated K18 proteins exhibited significantly perturbed tubulin assembly. Phosphorylation at S356 slightly impaired the K18-induced microtubule polymerization, whereas the di- and tri- phosphorylated forms of K18 bearing phosphorylation at S262 and S356 or S258, S262, and S356 strongly reduced microtubule polymerization. These findings suggest that phosphorylation at S262 has a dominant effect on the disruption of tau binding to microtubules.

**Fig. 5.**
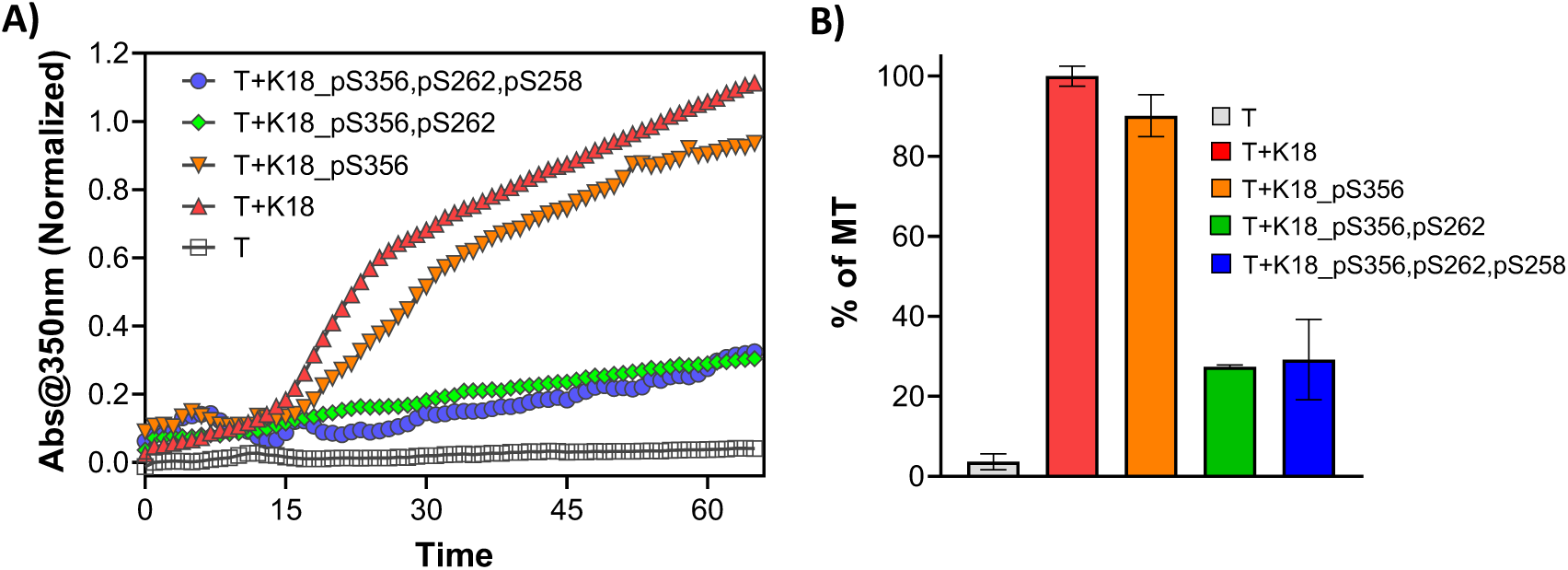
K18 proteins binding to microtubule. A) MT assembly in the presence of K18 proteins (K18, K18_pS356, K18_pS356, pS262 and K18_pS356, pS262, pS258) was evaluated by measuring light scattering at 350 nm over time. B) The percent of MT formed after 60 min of incubation with K18 proteins determined by measuring the absorbance at 350 nm (3 repeats, represented as the mean ± SD).

## Discussion

Despite the consistent observation that pathological tau species in the brains of patients with AD and other tauopathies are hyperphosphorylated, the role of tau phosphorylation in tau pathology formation and the pathogenesis of AD remains unclear. For decades, the prevailing hypothesis has been that hyperphosphorylation of tau induces its disassociation from microtubules, disrupts axonal transport and increases aggregation into PHFs. Testing this hypothesis has proven to be challenging for several reasons: 1) The exact residues and patterns of phosphorylation that define each hyperphosphorylated state remain poorly defined, especially since the state of phosphorylation is defined primarily using a limited set of antibodies, and the impact of the presence of multiple PTMs on the detection by these antibodies has not been systematically investigated; 2) The lack of synthetic strategies that enable the site-specific introduction of PTMs and preparation of homogeneously modified forms of tau has led experimentalists to rely primarily on pseudophosphorylated tau (S/T àD/E mutants) produced in *E. coli*.; and 3) *In vitro* phosphorylation using kinases has been used to generate hyperphosphorylated forms of tau, but this approach is not specific and leads to the generation of complex mixtures of tau phosphorylated species, thus complicating the analysis of the data and precluding an accurate assessment of the relative contribution of the different phosphorylation sites.

To address these limitations and pave the way for deciphering the tau PTM code, we developed and optimized a total chemical synthesis method for the K18 fragment and used it to generate K18 fragments that were site-specifically phosphorylated at single or multiple physiologically and pathologically relevant sites. The flexibility of this method allowed us for the first time to begin to answer with great precision the long-standing question of whether phosphorylation of tau within the microtubule domain inhibits or accelerates its aggregation. In addition, the availability of these homogeneously phosphorylated proteins enabled us to identify which phosphorylation sites dominantly regulate tau microtubule binding and aggregation, thus providing critical insight for developing novel therapeutic strategies on the basis of site-specific regulation of tau phosphorylation states.

Our results demonstrate that the effect of phosphorylation on K18 aggregation and K18 fibril seeding activity is sequence context-dependent and is dramatically reduced with an increasing number of phosphorylation sites. We showed that phosphorylation of K18 within the MTBD strongly inhibits fibril formation *in vitro* and leads to the accumulation of nonfibrillar oligomers. The reduced seeding activity for the triphosphorylated K18 suggests that the phosphorylated oligomers exhibited very low seeding activity. Our findings are consistent with previous studies by Mandelkow and colleagues demonstrating that *in vitro* hyperphosphorylation of tau by MARK and PKA kinases, which phosphorylate tau at multiple residues including S262 and S356, inhibits its aggregation^26^. However, these kinases phosphorylate tau at several additional sites, thus making it difficult to correlate the inhibitory effects observed with specific phosphorylation patterns^17^. In our work, we addressed this major limitation using protein total chemical synthesis, which enabled the generation of site-specifically and homogeneously hyperphosphorylated forms of K18.

In an attempt to gain insight into the structural basis underlying the inhibitory effects of phosphorylation and the relative contribution of each phosphorylation site, we reviewed the recent high-resolution atomic Cryo-EM structures of the tau filaments derived from the brains of patients with AD and Pick’s disease^12, 13^. These structures revealed that tau filaments are composed of C-shaped subunits with resolved R3 and R4 repeat regions (306–378 residues) of the MTBD (Fig. 6A). Interestingly, the high-resolution structure for the NPFs from Pick’s disease patient exhibits different molecular interactions and conformations than the PHFs from AD patients. The minimal effect of phosphorylation at S356 on K18 fibrillization and seeding activity could be explained by the fact that the side chain of S356 does not participate in any major stabilizing interactions in either PHF. Similarly, in the context of the NPFs, S356 is not part of the sequence that constitutes the core of the filaments; it is located in the unstructured region (^354^IGSLD^358^) and does not seem to participate in major interactions with other key residues involved in stabilizing the filament formation. In contrast, the side chains of serine residues S262 and S258 exhibit spatial hydrogen bonding with adjacent AA residues in NFTs (Fig. 6A). The S262 and S258 residues form hydrogen bonds with D264 and H362, respectively. Thus, phosphorylation of S262 would prevent hydrogen bonding with E264 and significantly affect the molecular interaction between K259 and D264 through electrostatic repulsion. Similarly, phosphorylation of S258 would prevent hydrogen bonding with H362. These structural attributes clearly imply that phosphorylation at S262 and S258 could cause significant structural perturbations that could affect protein folding and aggregation. This could possibly explain the drastic decrease in the aggregation propensity of the di- (K18_pS356, K18_pS262) and tri- (K18_pS356, K18_pS262, K18_pS258) phosphorylated K18 proteins. The increase in the number of phosphorylation sites significantly perturbs key molecular interactions implicated in mediating pathological folding, leading to the complete inhibition of fibril formation. This clearly indicates the cumulative effect of the three phosphorylation sites on the inhibition of fibrillization. It is noteworthy that despite the fact that the R1 repeat is not part of the core of ex vivo filaments^39^, several studies have consistently shown that sequence motifs within these region associated with disease-associated mutations^40^, PTMs^41^ or other sequence perturbations significantly alter the kinetics and aggregation properties of tau. Our results show that phosphorylation within the MTBD inhibits tau fibril formation, which is contrary to the conventional wisdom that this enhances tau aggregation and fibrillization.

**Fig. 6.**
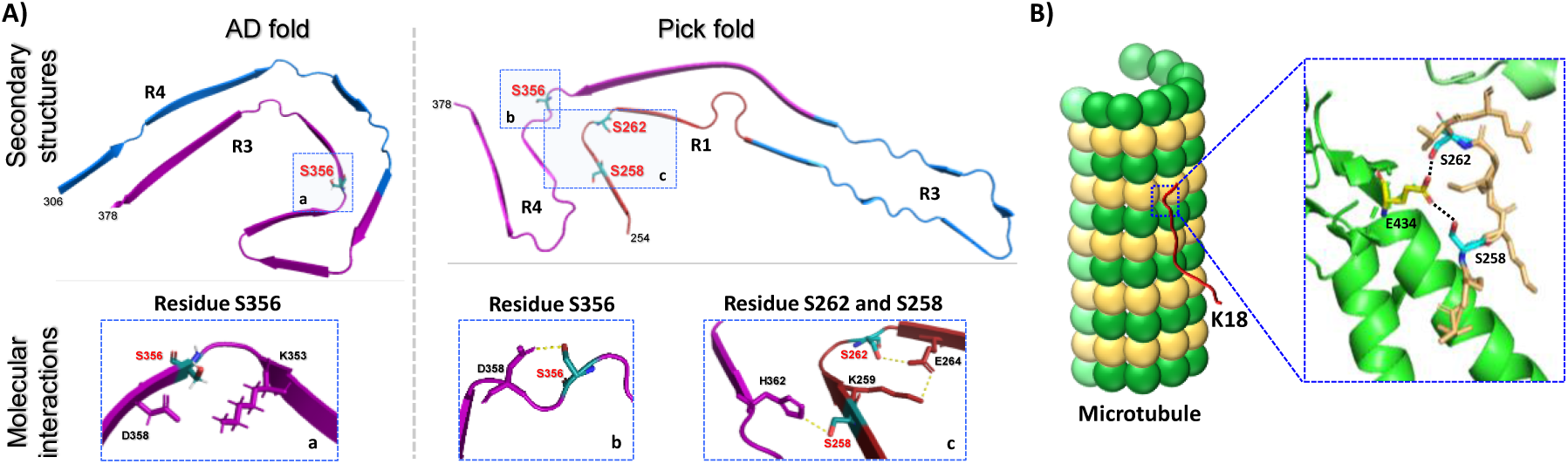
A) Rendered view of the secondary structure elements observed in Alzheimer’s and Pick’s filaments derived from the brains of AD and Pick’s disease patients. The repeats R1, R3, and R4 of MTBD are highlighted in red, blue and purple, respectively. The polar interactions (hydrogen bonding) for the side chains of residues S356, S262 and S258 identified in AD and Pick’s disease fold. B) Pictorial representation of the K18 fragment interacting with MT (left) and atomic model for the molecular interaction of pS258 and pS262 with E434 of the microtubule. The Cryo-EM structure reveals that S258, similar to S262, is in close proximity to MT and interacts with the E434 residue of MT, indicating possible roles of S258 and S262 in stabilizing the tau-MT complex.

Previous models that sought to establish a link between the disruption of tau normal function (stabilization of microtubules) and its propensity to aggregate speculated that the phosphorylation events responsible for inducing tau disassociation from microtubules are likely to also increase aggregation and pathology formation. However, the lack of methods for the site-specific phosphorylation of tau made it difficult to evaluate and test these models. Our work here shows this is not the case. For example, phosphorylation at S356, which does not influence K18 aggregation and seeding, results in small but significant inhibition of tau-mediated microtubule assembly. On the other hand, phosphorylation of the S262 residue, which results in a dramatic reduction of microtubule polymerization, resulted in significant inhibition of tau fibril formation and cellular seeding activity. Interestingly, phosphorylation at both S258 and S262 resulted in the near abolishment of K18 fibrillization *in vitro*, seeding activity in cells and tau-mediated microtubule polymerization. These results show that phosphorylation events within the MTBD that lead to the disruption of tau microtubule binding also inhibit its aggregation and seeding activity. These findings are consistent with previous studies showing that the nonspecific phosphorylation of tau using kinases that phosphorylated at S262 or the introduction of point mutations that mimic phosphorylation at this residue disrupt tau binding to microtubules and inhibit tau-mediated tubulin assembly^42^. Together, these observations suggest that S262 plays a key role in the tubulin-tau interaction and the stabilization of this native conformation of the protein. This is consistent with a recent report on the atomic structure of tubulin and tau interactions, which shows that both residues S262 and S258 of tau are directly involved in hydrogen bonding interactions with tubulin residue E434 (Fig. 6B), which further highlights the role of S262 in MT binding.^43^ Introducing phosphorylation at S262 and S258 would disrupt the hydrogen bonding and further introduce electrostatic repulsion between pS262 or pS258 of tau and tubulin E434, which could potentially destabilize the tau-MT interaction. These attributes clearly support our experimental results that K18 phosphorylated at S262 and S258 significantly disrupts MT assembly.

### Implications for tau therapeutic strategies

Disassociation of tau from MTs has been proposed as one of the key early pathological events that lead to increasing the pool of free tau, which then aggregates and forms PHFs^7^. Therefore, stabilizing tau interactions with microtubules or interfering with events that lead to its disassociation from microtubules (i.e., native state stabilization) could constitute an effective strategy to prevent tau aggregation and pathology formation. On the basis of our results and previous findings discussed above, we propose that inhibiting the kinases responsible for the phosphorylation events that lead to the disruption of the tau interaction with microtubules represents an attractive strategy for preventing tau accumulation and aggregation via stabilizing its native microtubule-bound conformation. Given the dominant role of S262 phosphorylation in disrupting tau interactions with MTs, we propose that inhibiting the natural kinases responsible for phosphorylating tau at S262 or S258 and S262 constitutes a viable therapeutic strategy for the treatment of AD and potentially other tauopathies. This would not only stabilize the native state of the tau–Tubulin complex (Fig. 7) but would also decrease the levels of unbound tau available for the formation of potentially toxic species (oligomers and fibrils). As phosphorylation at S262 alone had only modest effect on tau aggregation compared to the di- and triphosphorylated K18 protein, we do not anticipate that selective inhibition of phosphorylation at this residue would promote tau aggregation.

**Fig. 7.**
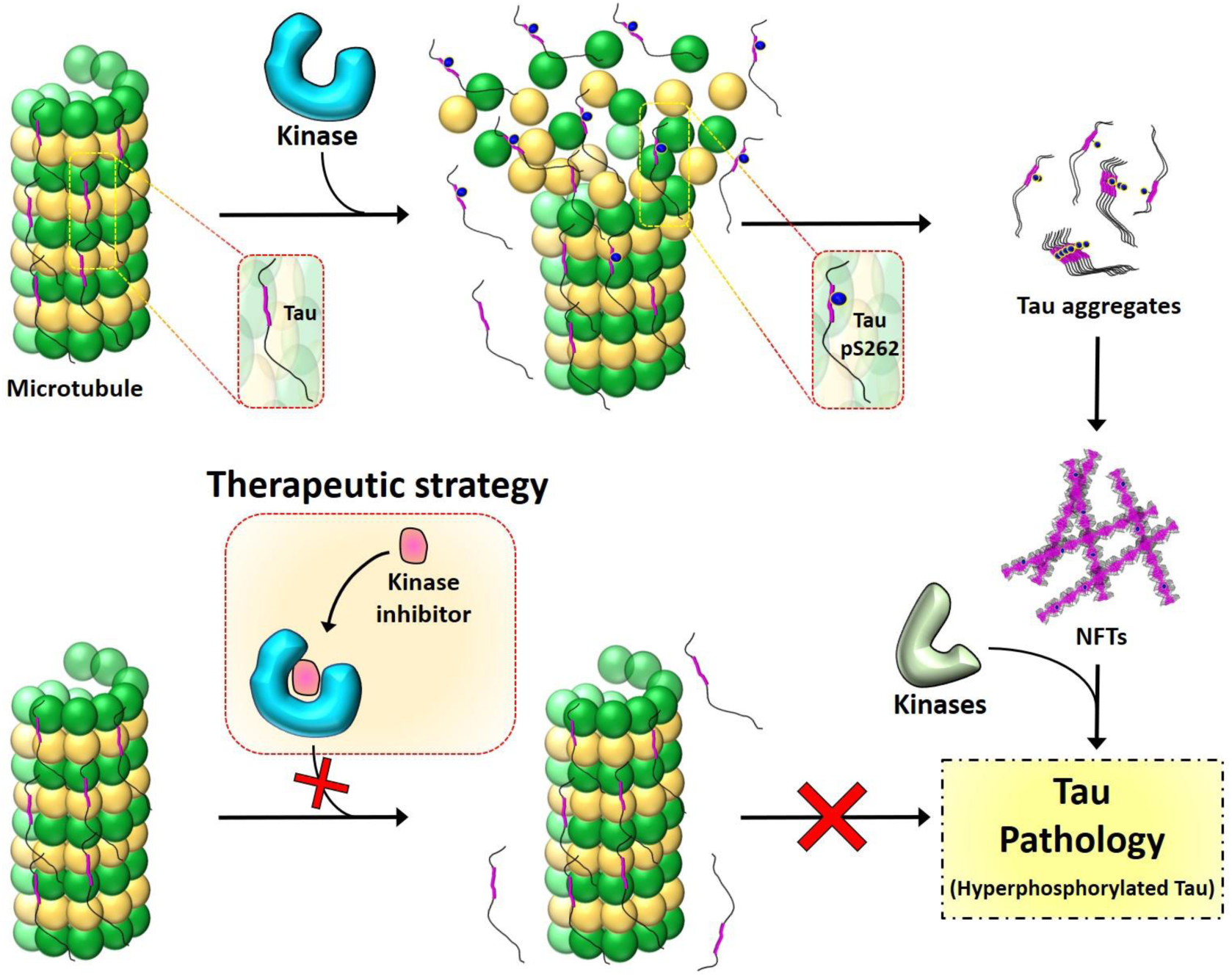
The residue S262 plays a vital role in stabilizing the binding of tau to tubulin of MTs; therefore, phosphorylation at S262 adversely affects the tau-microtubule interaction (top), subsequently leading to tau pathology. Inhibiting the phosphorylation at S262 by targeting the kinases involved could be an effective therapeutic strategy (bottom) to prevent tau dissociation from MTs and stabilize the native state of the tau–Tubulin complex.

Several kinases have been identified to phosphorylate the MTBD; however, they lack selectivity and phosphorylate at multiple sites^25^. For example, kinases such as glycogen synthase kinase 3 (GSK3) and casein kinase 1\2 (CK1\2) extensively phosphorylate the MTBD at multiple residues, including S258 and S262^44^. Similarly, AMP-activated protein kinase (AMPK) predominantly phosphorylates KXGS repeat units of the MTBD, which also includes the residues S258 and S262^45^. Interestingly, the levels of GSK3β^46^, CK1^47^ and AMPK^48^ are increased several-fold in AD. Moreover, these kinases predominantly colocalize with the NFTs observed in the brains of patients with AD.

Microtubule affinity-regulating kinase 1 (MARK1), salt-inducible kinase (SIK), cyclic AMP-dependent protein kinase (cAMP-PK) and calcium/calmodulin-dependent protein kinase II (CaMKII) mostly phosphorylate residues S262 and S356 of the MTBD^49, 50^. Interestingly, kinases such as checkpoint kinase 1/2 (Chk 1/2)^51^, P70 S6^52^ and phosphorylase b kinase (PhK)^53^ phosphorylate only S262 among all AD relevant phosphorylating sites (S258, S262, S289 and S356) in the MTBD. Similarly, protein kinase N (PKN) and protein kinase C (PKC) preferentially phosphorylate at the S258 residue^54^. We propose that inhibiting the activity of these kinases or others that preferentially target these residues could lead to native state stabilization of tau and thus prevent its aggregation and pathology formation. Studies are underway to evaluate this hypothesis and therapeutic approach. In addition, we plan to map the different phosphorylation events that directly or indirectly disrupt tau MT binding with the aim of expanding the range of possible target kinases and kinase inhibitors as potential modulators of tau aggregation, toxicity and pathology spreading in cellular and animal models of tauopathies.

## Conclusions

Taken together, our results argue against the prevailing hypothesis that phosphorylation promotes tau aggregation and pathology formation. Instead, our data show that phosphorylation within the MTBD inhibits tau aggregation and seeding activity in cells with the inhibitory effects increasing with the increasing number of phosphorylated sites, with phosphorylation at S258 and S262 having a dominant effect on K18 aggregation. The fact that the level of phosphorylation at these sites has been found to be elevated in the pathology of AD and other tauopathies may reflect post aggregation phosphorylation events rather than the primary role of phosphorylation at these residues in driving pathology formation.

Together, our findings highlight the potential of targeting tau phosphorylation for the treatment of AD and tauopathies and underscore the critical importance of revisiting the role of site-specific phosphorylation and hyperphosphorylation in regulating tau interactions with microtubules and binding partners with tau and transcellular pathology propagation. Furthermore, our results demonstrate that the oligomerization, fibrillization and tubulin binding properties of K18 are highly sensitive to its phosphorylation state and pattern. Thus, underscoring the importance of reassessing the phosphorylation patterns associated with the different physiological and pathogenic hyperphosphorylated states of tau with an emphasis on determining which phosphorylation sites cooccur on the same molecule. This, combined with the advances made by our group^8^ and others^55, 56^ to enable site-specific modifications of tau, should enable the generation and characterization of the different, modified tau species in homogeneous forms, thus paving the way for more systematic investigations of the tau phosphorylation code in health and disease. Our data also provide strong evidence in support of pursuing native state stabilization strategies for the treatment of AD and tauopathies. It is noteworthy that the only FDA-approved small molecule aggregation inhibitor is one that acts by stabilizing the native tetrameric state of the amyloidogenic protein Transthyretin^57^. While tau does not have a stable folded monomeric conformation in solution or ligand binding pockets that could be targeted by small molecules, it does engage in highly specific and stable molecular interactions where it becomes highly ordered^58^. Developing and identifying strategies for stabilizing these interactions through the modulation of phosphorylation or other modifications could provide viable small molecule-based strategies for targeting tau aggregation and toxicity. A recent study by Sharma et al. showed that monomeric tau exists in multiple conformations that possess distinct seeding activity^59^. These finding also imply that stabilizing the native seeding incompetent tau monomers could also provide an alternative mechanism for blocking aSyn aggregation, seeding and pathology formation. However, further studies are needed to determine the role of phosphorylation at different residues in regulating the equilibrium between seeding competent and seeding incompetent monomers.

## Acknowledgments

This work was supported by the École Polytechnique Fedéralé de Lausanne. We thank Dr. Nadine Ait Bouziad for her feedback and review of the manuscript.

## Author Information

Pushparathinam Gopinath: Present address - Department of Chemistry, SRM Institute of Science and Technology, Chennai, Tamilnadu, India.

## Author contributions

H.A.L. and M.H.Y. conceived the project. M.H.Y. designed the total synthesis, M.H.Y. and P.G. synthesised the proteins and preformed the in vitro aggregation studies. M.I.D. and H.M. performed seeding studied in biosensor cells and analysed the data. K.R. performed microtubule polymerisation assay. H.A.L., M.H.Y., and K.R. analysed the results. H.A.L. and M.H.Y. and K.R took the lead in writing the manuscript. P.G., M.I.D., H.M., and K.R. contributed in writing their respective experimental results.

## Supplementary Information

### Synthesis of tau(322-372)-CONH2 (1)

**Sequence:**CGSLGNIHHKPGGGQVEVKSEKLDFKDRVQSKIGSLDNITHVPGG GNKKIE **tau (322-372) (1)** was synthesized using a Rink amide-MBHA resin (0.27 mmol/g, 0.1 mmol scale) according to the following procedure: the first amino acid (AA), glutamine (Glu), was double coupled manually for 1 h (×2) using 4 eq of 1-[bis(dimethylamino)methylene]-1H-1,2,3-triazolo[4,5-b]pyridinium3-oxid hexafluorophosphate (HATU) and 8 eq of N,N-diisopropylethylamine (DIEA). The remaining AAs were double coupled using an automated CS 336X peptide synthesizer from CS Bio and carried out by mixing 4 eq of the AA with 8 eq of DIEA and 4 eq of O-(1H-6-chlorobenzotriazole-1-yl)-1,1,3,3-tetramethyluronium hexafluorophosphate (HCTU) in N-methyl 2-pyrrolidone (NMP). Each coupling reacted for 45 min, and Fmoc deprotection was achieved using 20% piperidine and 50 mM 1-hydroxybenzotriazole hydrate (HOBt) in 3/5/3 min cycles. The last AA, cysteine (Cys), was introduced with N-terminal Boc protection.

*Cleavage from the resin:* Once the synthesis was complete, the resin was washed with dimethylformamide (DMF), methanol, and DCM, and the dried peptidyl-resin was treated with a cleavage cocktail containing 95% trifluoroacetic acid (TFA), 2.5% water, and 2.5% triisopropylsilane (TIPS) for 2.5 h at room temperature (RT). The crude peptide was then precipitated by the dropwise addition of a 10-fold volume of cold ether and centrifuged. The pellet was dissolved in 30-50% aqueous acetonitrile and lyophilized. The crude peptide was then purified by RP-HPLC using a preparative C18 column with a linear gradient of 30-60% B over 40 min, resulting in a 25% yield of the final peptide (Fig. S1).

**Fig. S1.**
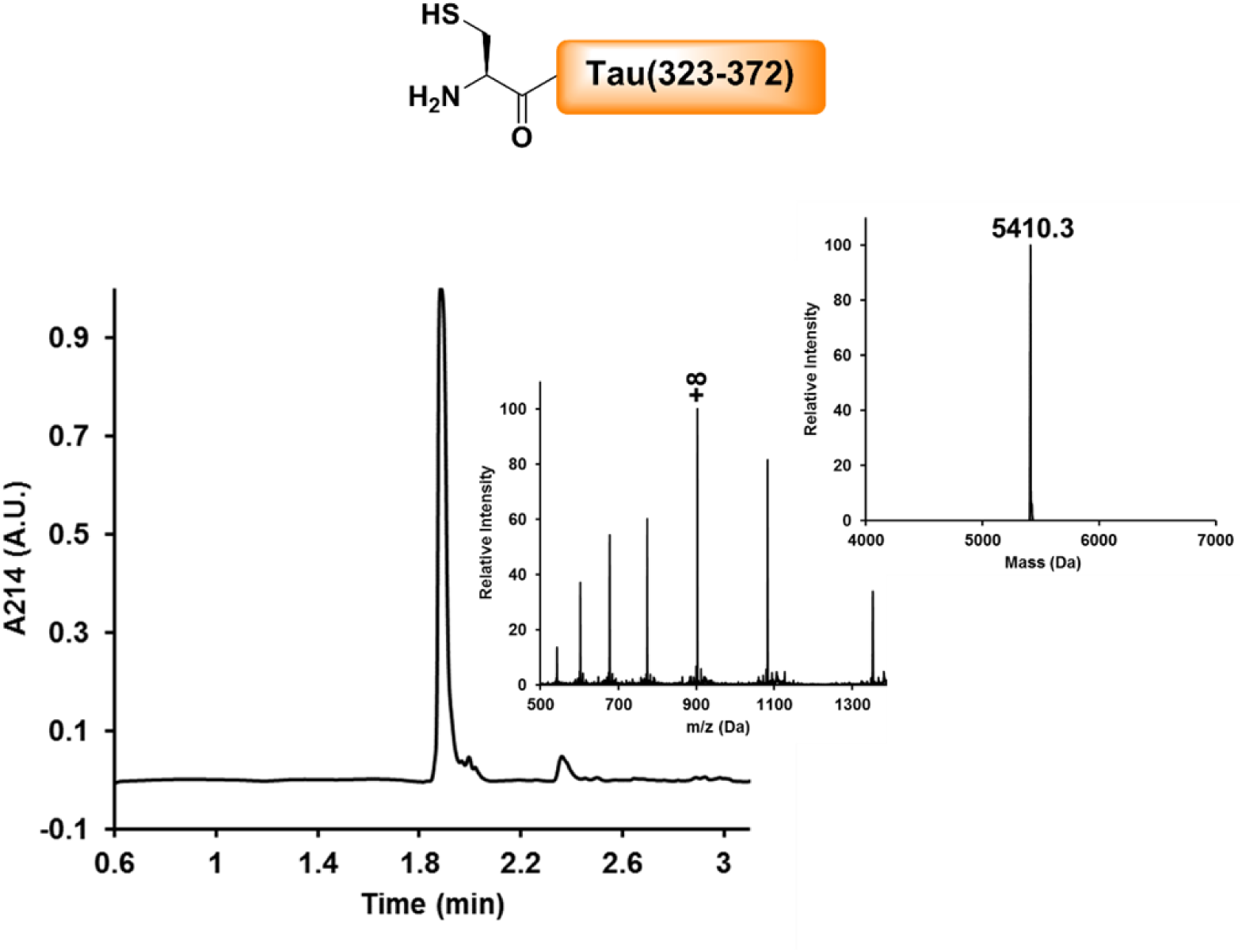
SPPS of tau (322-372). Analytical RP-UHPLC and ESI-MS of the purified peptide **1**, with the observed mass of 5410.3 Da (calculated 5409.13 Da).

### Synthesis of tau(322-372, pS356)-CONH2 (1a)

**Sequence:**CGSLGNIHHKPGGGQVEVKSEKLDFKDRVQSKIGpSLDN<u>I</u> <u>T</u>HVPGGGNKKIE **tau (322-272, pS356)-CONH2 (1a)** was prepared as described for peptide **1**, with the following modifications: 2.5 eq of Fmoc-Ser(HPO3Bzl)-OH was coupled at position 356 for 2 h. Additionally, 2.5 eq of the pseudoproline dipeptide of Ile-Thr were manually coupled for 2 h at position Ile360-Thr361. The lyophilized crude peptide was then purified by RP-HPLC using a preparative C18 column with a linear gradient of 30-60% B over 40 min, resulting in ∼20% yield of the final peptide (Fig. S2).

**Fig. S2.**
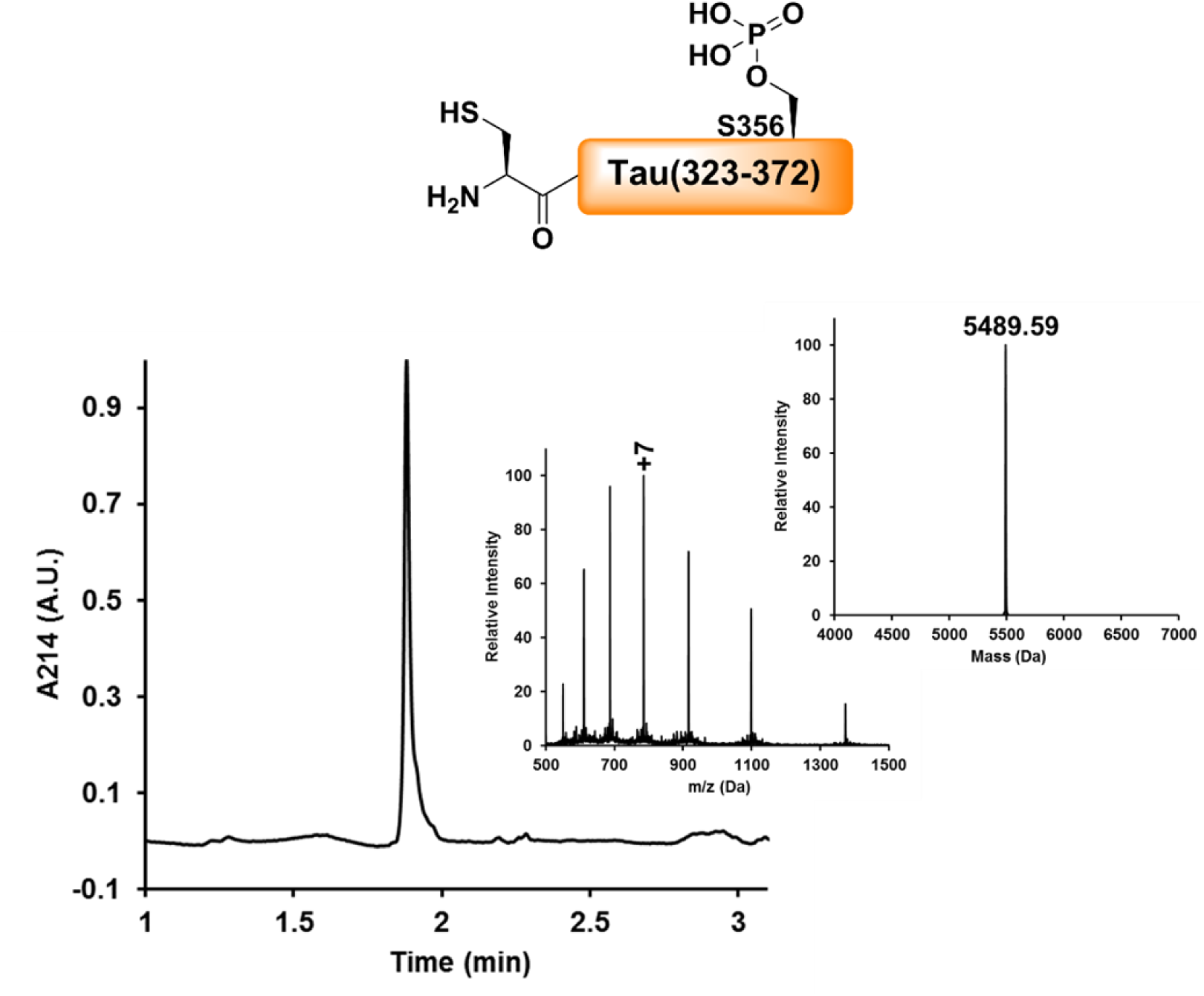
SPPS of tau (322-372, pS356). Analytical RP-HPLC and ESI-MS of the purified peptide **1a**, with the observed mass of 5489.59 Da (calculated 5489.10 Da).

### Synthesis of tau (C291Thz-321)-SR (2)

Sequence: **Thz-GSKDNIKHVPGGGSVQIVYKPVDLSKVTSK tau (C291Thz-321)-SR (2)** was synthesized using Rink amide-MBHA resin (0.27 mmol/g, 0.1 mmol scale) according to the following procedure: The first unnatural AA, 3-(Fmoc-amino)-4-(methylamino) benzoic acid, was double coupled manually for 1 h (×2) using 4 eq of HATU and 8 eq of DIEA. The remaining AAs were double coupled using an automated CS 336X peptide synthesizer from CS Bio and carried out in the presence of 4 eq of AA, 8 eq of DIEA and 4 eq of HCTU. Each coupling reacted for 45 min, and Fmoc deprotection was achieved using 20% piperidine and 50 mM HOBt in 3/5/3 min cycles. The last AA was introduced with N-terminal Boc protection. After peptide elongation, the resin was reswelled with dichloromethane (DCM) for 15 min and reacted with a solution of p-nitrophenyl chloroformate (5 eq in 2 ml of DCM) for 1 h at RT. Then, the resin was washed with DCM and reswelled with DMF for 15 min. To that, a solution of 0.5 M DIEA in 1 ml of DMF was added for 15 min (×2).

*Cleavage from the resin:* After washing with DMF, methanol, and DCM, the dried peptidyl-resin was treated with a cleavage cocktail containing 95% TFA, 2.5% water and 2.5% TIPS for 2.5 h at RT. The crude peptide was then precipitated by the dropwise addition of a 10-fold volume of cold ether and centrifuged. The precipitate was then dissolved in 30-50% aqueous acetonitrile and lyophilized. The crude peptide was then purified by RP-HPLC using a preparative C18 column with a linear gradient of 10-40% B over 40 min, resulting in 22.5% yield of the thioester (Fig. S3).

**Fig. S3.**
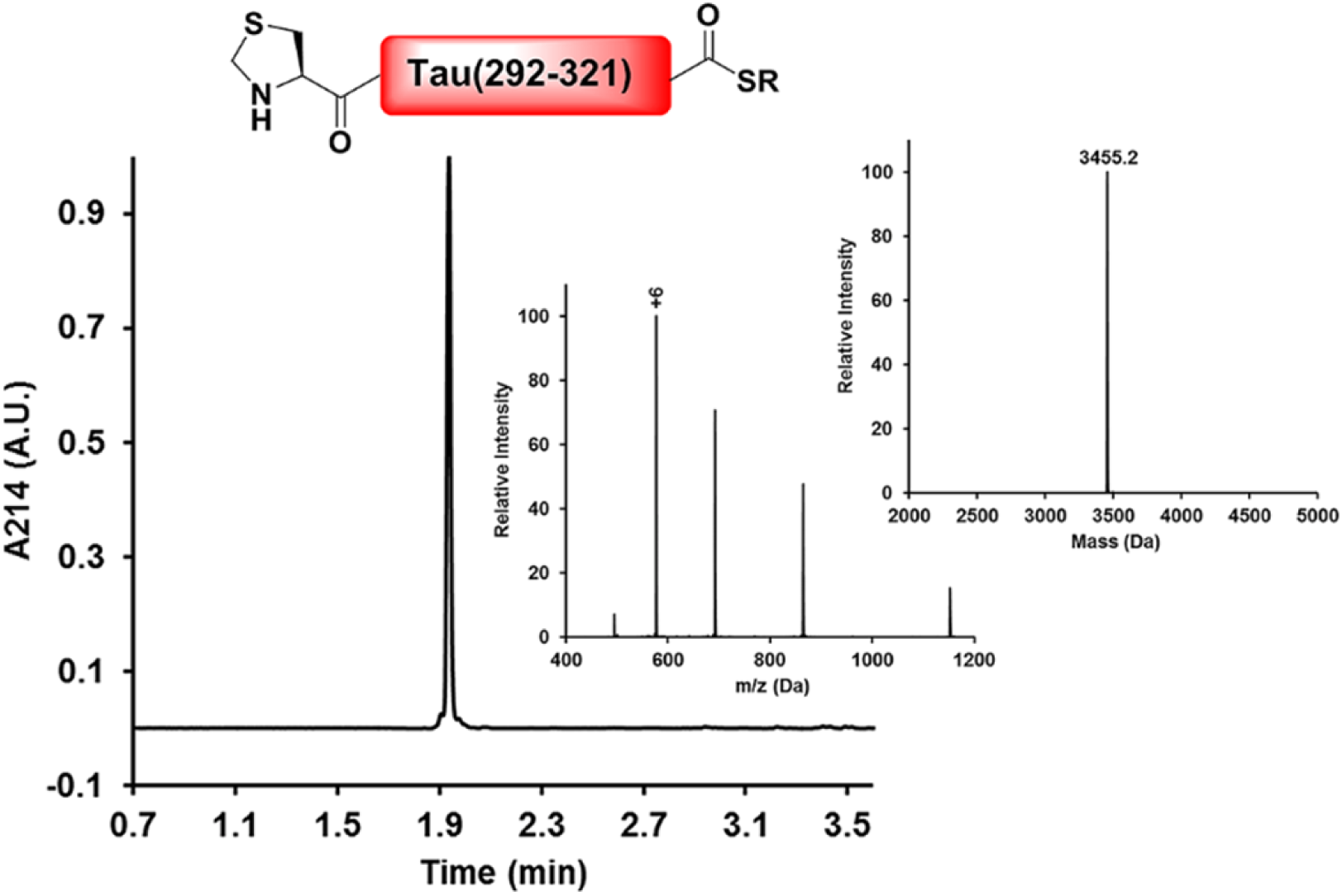
SPPS of tau (C291Thz-321)-SR. Analytical RP-HPLC and ESI-MS of the purified thioester **(2)**, with the observed mass of 3455.20 Da (calculated 3455.50 Da).

### Synthesis of tau (243-290)-SR (3)

Sequence: LQTAPVPMPDLKNVKSKIGSTENLKHQPGGGKVQIINKKLD<u>LS</u>NVQSK **tau (243-290)-SR (3)** was synthesized as described above with the following modifications: 2.5 eq of the pseudoproline dipeptides of Leu-Ser was manually coupled for 2 h at the Leu284-Ser285 junction. The peptide was purified by RP-HPLC using a preparative C18 column with a linear gradient of 30-60% B over 40 min, resulting in ∼15% yield of the thioester (Fig. S4).

**Fig. S4.**
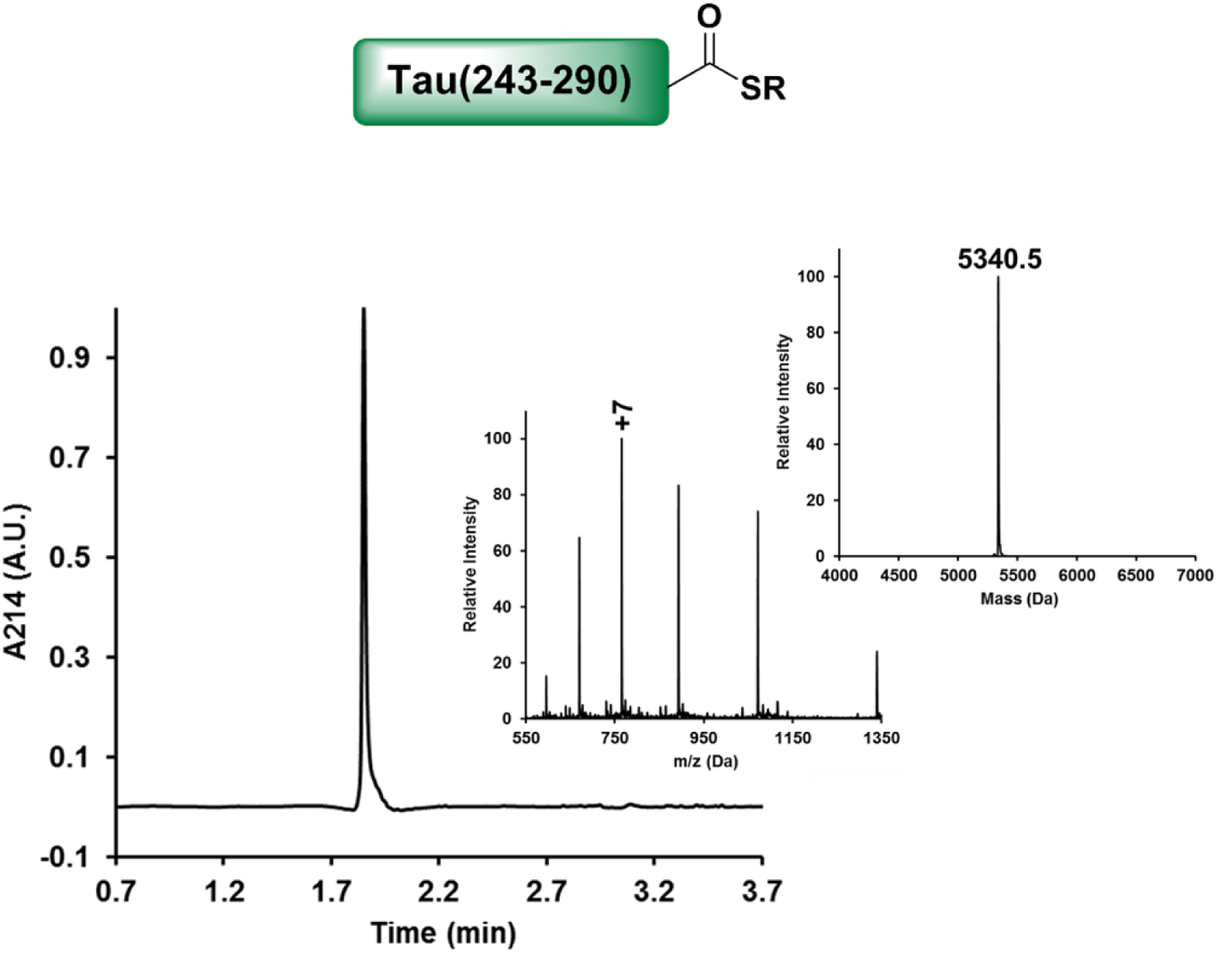
SPPS of tau (243-290)-SR. Analytical RP-HPLC and ESI-MS of the purified peptide **(3)**, with the observed mass of 5340.50 Da (calculated 5337.05 Da).

### Synthesis of tau(243-290, pS262)-SR (3a)

Sequence: LQTAPVPMPDLKNVKSKIGpSTENLKHQPGGGKVQIINKKLDLSNVQSK **tau (243-290, pS262)-SR (3a)** was synthesized similarly to the WT peptide **(3)** described above with the following modification: 2.5 eq of Fmoc-Ser(HPO3Bzl)-OH was coupled manually at position 262 for 2 h. The peptide was purified by RP-HPLC using a preparative C18 column with a linear gradient of 30-60% B over 40 min, resulting in an 8% yield of the thioester (Fig. S5).

**Fig. S5.**
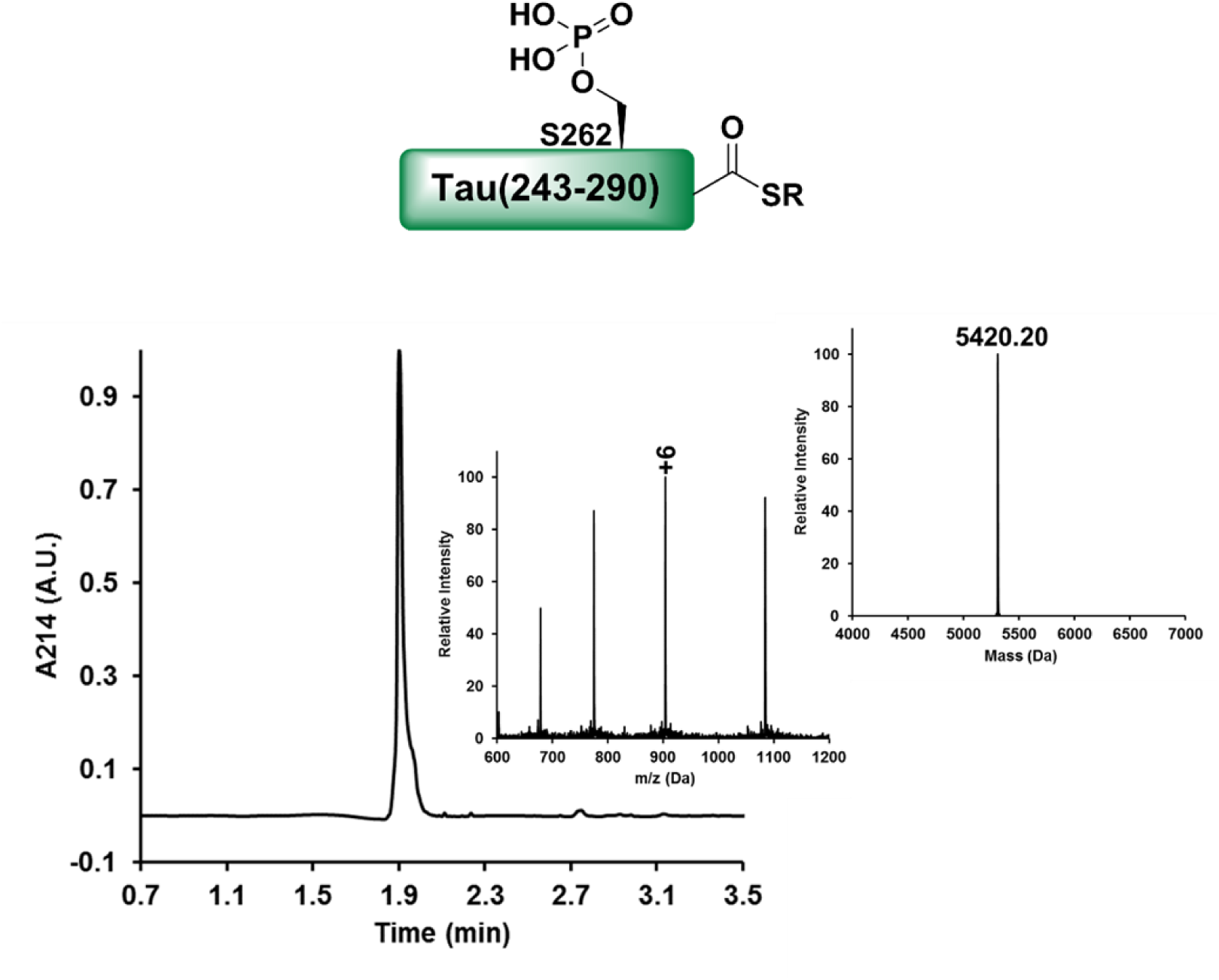
SPPS of tau (243-290, pS262)-SR. Analytical RP-HPLC and ESI-MS of the purified peptide **3a**, with the observed mass of 5420.20 Da (calculated 5417.05 Da).

### Synthesis of tau(243-290, pS262, pS258)-SR (3b)

Sequence: LQTAPVPMPDLKNVKpSKIGpSTENLKHQPGGGKVQIINKKLDLSNVQSK **tau (243-290, pS262, pS258)-SR (3b)** was synthesized similarly to peptide **3** described above with the following changes: 2.5 eq of Fmoc-Ser(HPO3Bzl)-OH was coupled manually at positions 262 and 258 for 2 h each. The peptide was purified by RP-HPLC using a preparative C18 column with a linear gradient of 30-60% B over 40 min, resulting in a 7.3% yield of the thioester (Fig. S6).

**Fig. S6.**
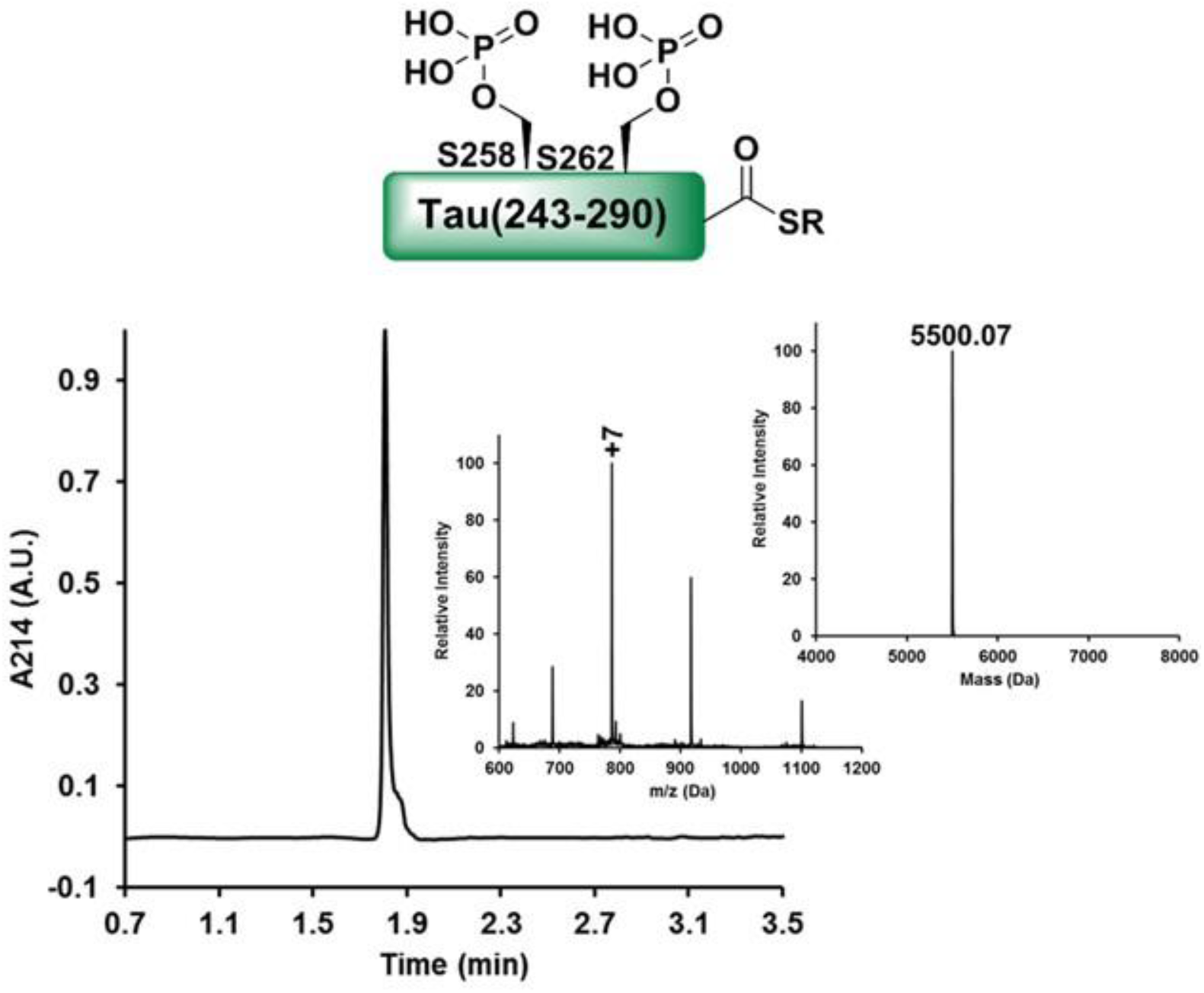
SPPS of tau (243-290, pS262, pS258)-SR. Analytical RP-HPLC and ESI-MS of the purified peptide **3b**, with the observed mass of 5500.07 Da (calculated 5497.05 Da).

### Ligation of tau (C291Thz-321)-SR with tau (322-372)

**tau (322-372) (1)** (10 mg, 1 eq) and **tau (C291Thz-321)-SR (2)** (9 mg, 1.4 eq) were dissolved in ligation buffer (6 M guanidine hydrochloride (GdnHCl), 0.2 M sodium phosphate, 25 eq tris(2-carboxyethyl)phosphine (TCEP), 20 eq 4-mercaptophenylacetic acid (MPAA), pH 7.0) that had been purged with nitrogen. The mixture was incubated at 37 °C for 4.5 h with orbital agitation at 600 rpm and monitored by RP-UHPLC and mass spectrometry analyses.

### In situ thiazolidine deprotection

Upon completion of the ligation reaction, a solution of 0.2 M methoxylamine, 30 eq TCEP in a nitrogen purged 6 M GdnHCl, 0.2 M phosphate was added to the ligation mixture and then incubated at 37 °C for 12 h with orbital agitation at 600 rpm. The reaction was monitored by RP-UHPLC and mass spectrometry analyses. The ligated product was purified by RP-HPLC using a semi-preparative C4 column with a linear gradient of 20% (30 min)-35% B over 50 min, resulting in 17.2% yield of the product **4** over two steps (Fig. S7).

**Fig. S7.**
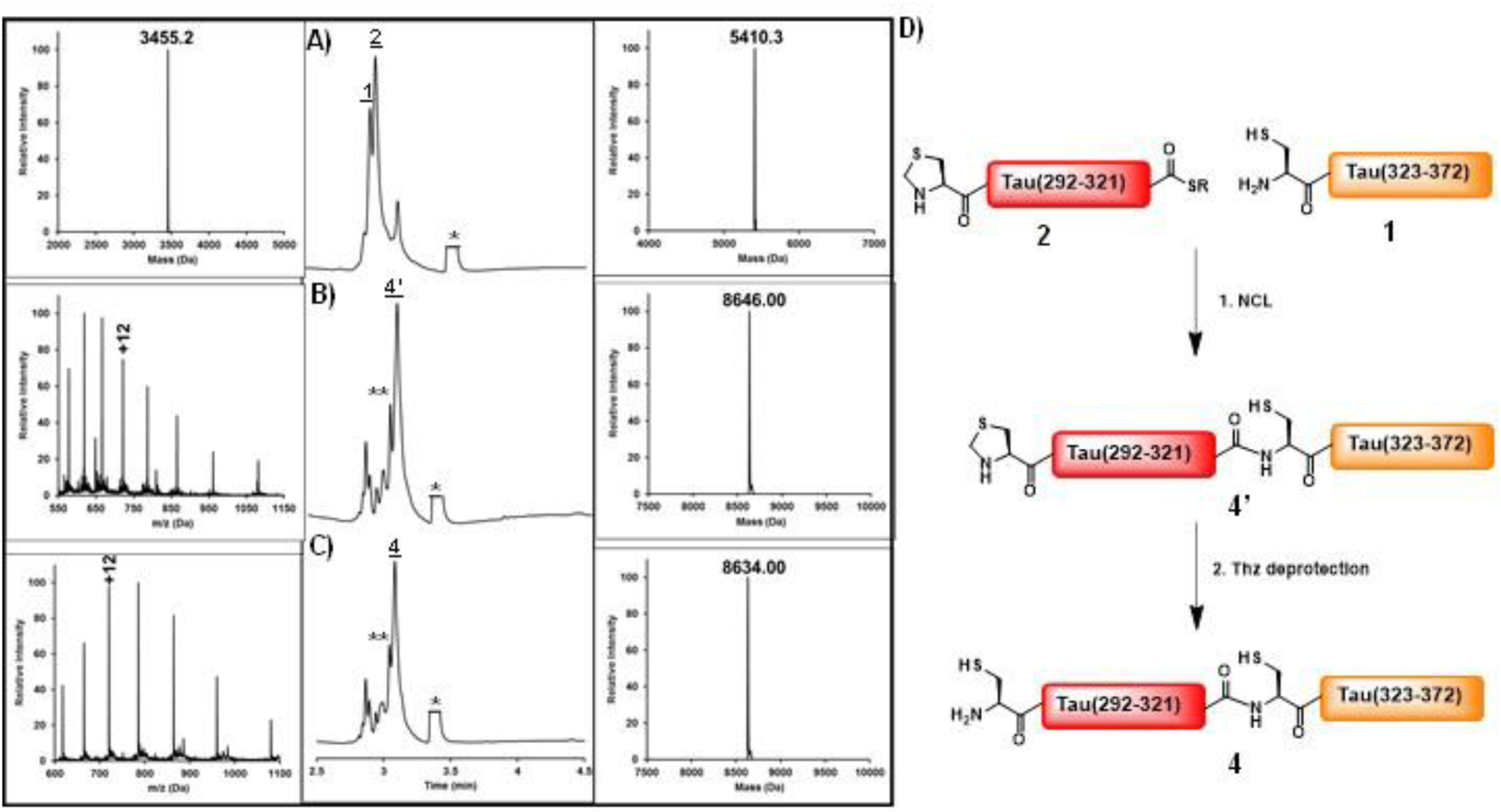
Native chemical ligation for the preparation of fragment 4. Analytical RP-HPLC and ESI-MS spectra of the ligation reaction between tau (322-372) and tau (C291Thz-321)-SR at A) 0 h and after B) 4.5 h. Peak **4** corresponds to the ligation product with the observed mass of 8646.00 Da (calculated 8646.00 Da). Peak ** corresponds to the hydrolyzed thioester, and peak * corresponds to the MPAA thiol. C) Analytical RP-HPLC and ESI-MS spectra of the in situ thiazolidine deprotection reaction after 12 h. Peak **4** corresponds to the product with free cysteine at the N-terminus, with the observed mass 8634.00 Da (calculated 8633.85 Da). D) Schematic representation of the NCL of the tau (291-372) fragment.

### Ligation of tau (243-290)-SR with tau (291-372)

**tau (291-372) (4)** (3 mg, 1 eq) and **tau (243-290)-SR (3)** (2.4 mg, 1.3 eq) were dissolved in NCL purged buffer (8.0 M urea, 25 mM TCEP, 2% TFET, pH 7.0). The mixture was incubated at 37 °C for 2 h with orbital agitation at 600 rpm and monitored by RP-UHPLC and mass spectrometry analyses. The ligated product was purified by RP-HPLC using a semipreparative C18 column with a linear gradient of 20-35% B over 50 min, resulting in 19% yield of the product **5** (Fig. S8).

**Fig. S8.**
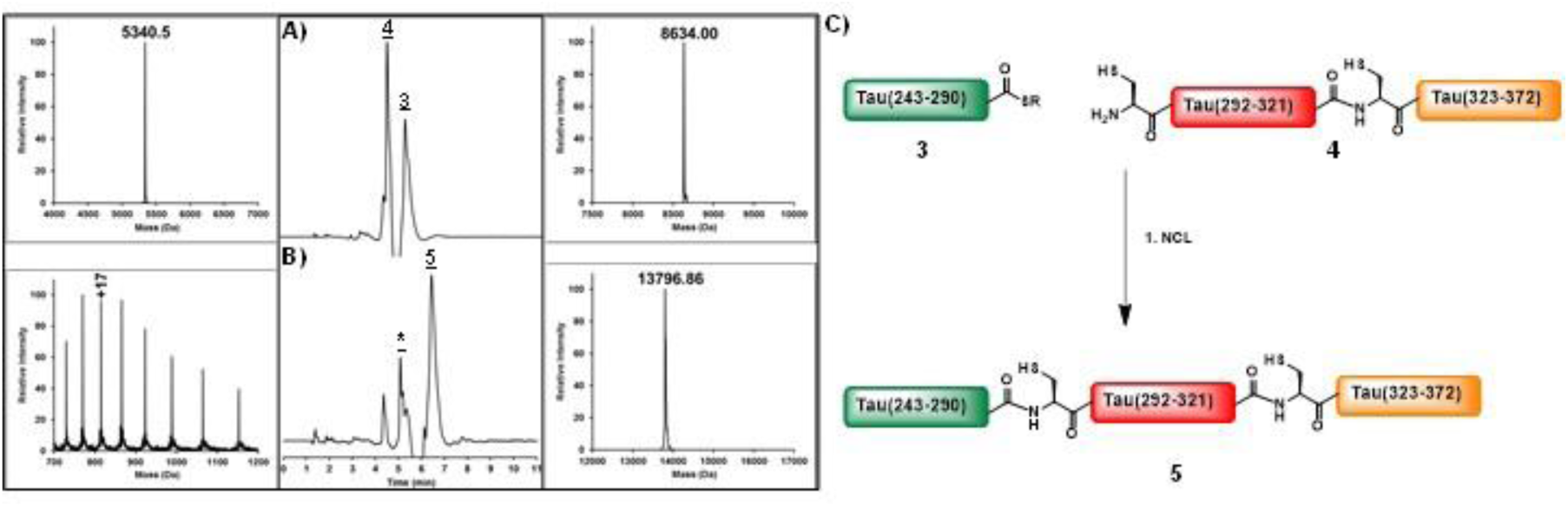
Native chemical ligation for the preparation of K18 (5) Analytical RP-HPLC and ESI-MS spectra of the ligation reaction between tau (243-290)-SR and tau (291-372) at A) 0 h and B) after 2 h. Peak **5** corresponds to the ligation product with the observed mass of 13796.00 Da (calculated 13795.00 Da). Peak * corresponds to the hydrolyzed thioester. C) Schematic representation of the total chemical synthesis of the tau (243-372) fragment (K18).

### Preparation of monophosphorylated K18 at Ser356

**tau (322-372, pS356) (1a)** (7 mg, 1 eq) and **tau (C291Thz-321)-SR (2)** (5.7 mg, 1.3 eq) were dissolved in ligation buffer (6.0 M GdnHCl, 25 mM TCEP, 2% TFET, pH 7.0). The mixture was left at 37 °C for 1.5 h with orbital agitation at 600 rpm and monitored by RP-UHPLC and mass spectrometry.

### In situ thiazolidine deprotection

Upon completion of the ligation reaction, a solution of 0.2 M methoxylamine and 30 eq of TCEP in a nitrogen purged 6.0 M GdnHCl, 0.2 M phosphate solution was added to the ligation mixture and then incubated at 37 °C for 12 h with orbital agitation at 600 rpm. The reaction was monitored by RP-UHPLC and mass spectrometry analyses. The ligated product was purified by RP-HPLC using a semi-preparative C18 column with a linear gradient of 20-35% B over 50 min, resulting in 17.2% yield of product **4a** over two steps (Fig. S9).

**Fig. S9.**
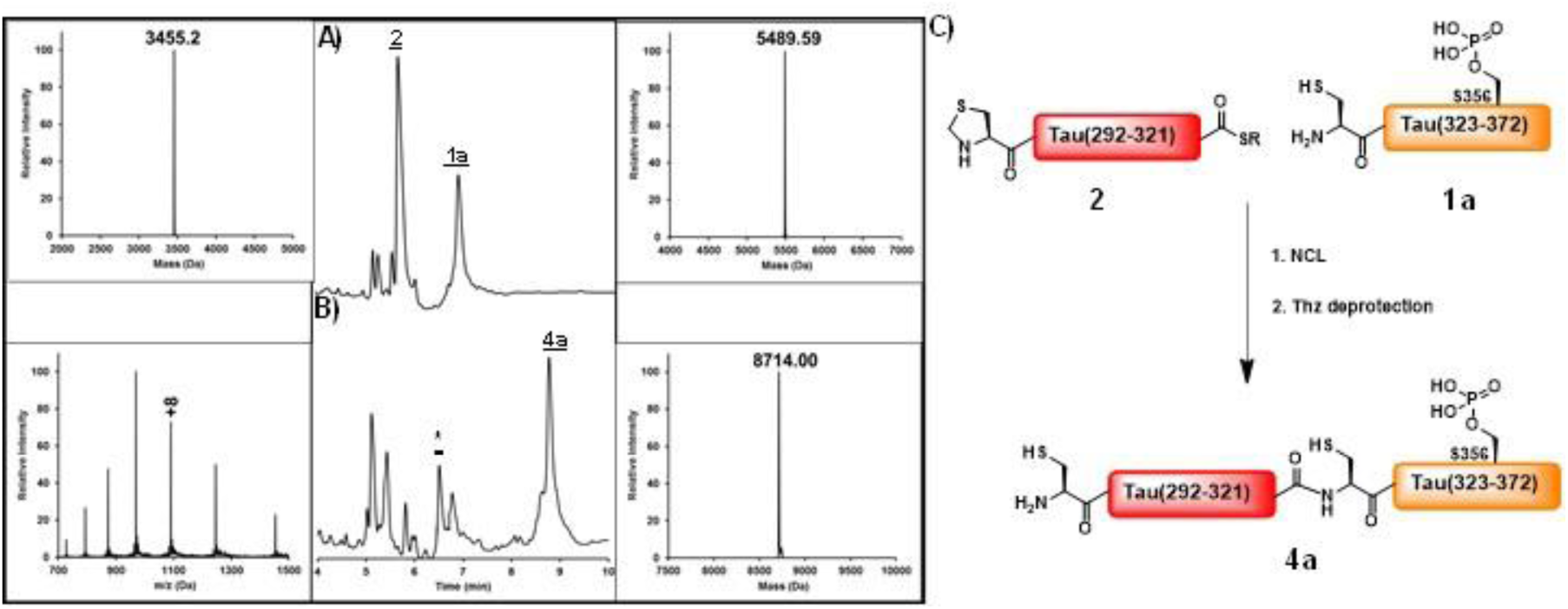
Native chemical ligation for the preparation of fragment 4a. Analytical RP-HPLC and ESI-MS spectra of the ligation reaction between tau (322-372, pS356) and tau (C291Thz-321)-SR at A) 0 h and B) after 1.5 h. Peak **4a** corresponds to the ligation product after Thz deprotection with the observed mass of 8714.00 Da (calculated 8713.85 Da). Peak * corresponds to the hydrolyzed thioester. C) Schematic representation of the NCL of the tau (291-372, pS356) fragment.

### Ligation of tau(243-290)-SR with tau(291-372, pS356)

**tau (291-372, pS356) (4a)** (2.8 mg, 1 eq) and **tau (243-290)-SR (3)** (2.2 mg, 1.3 eq) were dissolved in NCL purged buffer (6.0 M GdnHCl, 25 eq TCEP, 20 eq MPAA, pH 7.0). The mixture was incubated at 37 °C for 2 h with orbital agitation at 600 rpm and monitored by RP-UHPLC and mass spectrometry analyses. The ligated product was purified by RP-HPLC using a semipreparative C4 column with a linear gradient of 10-55% B over 55 min to afford the corresponding product **6** in 25% yield (Fig. S10).

**Fig. S10.**
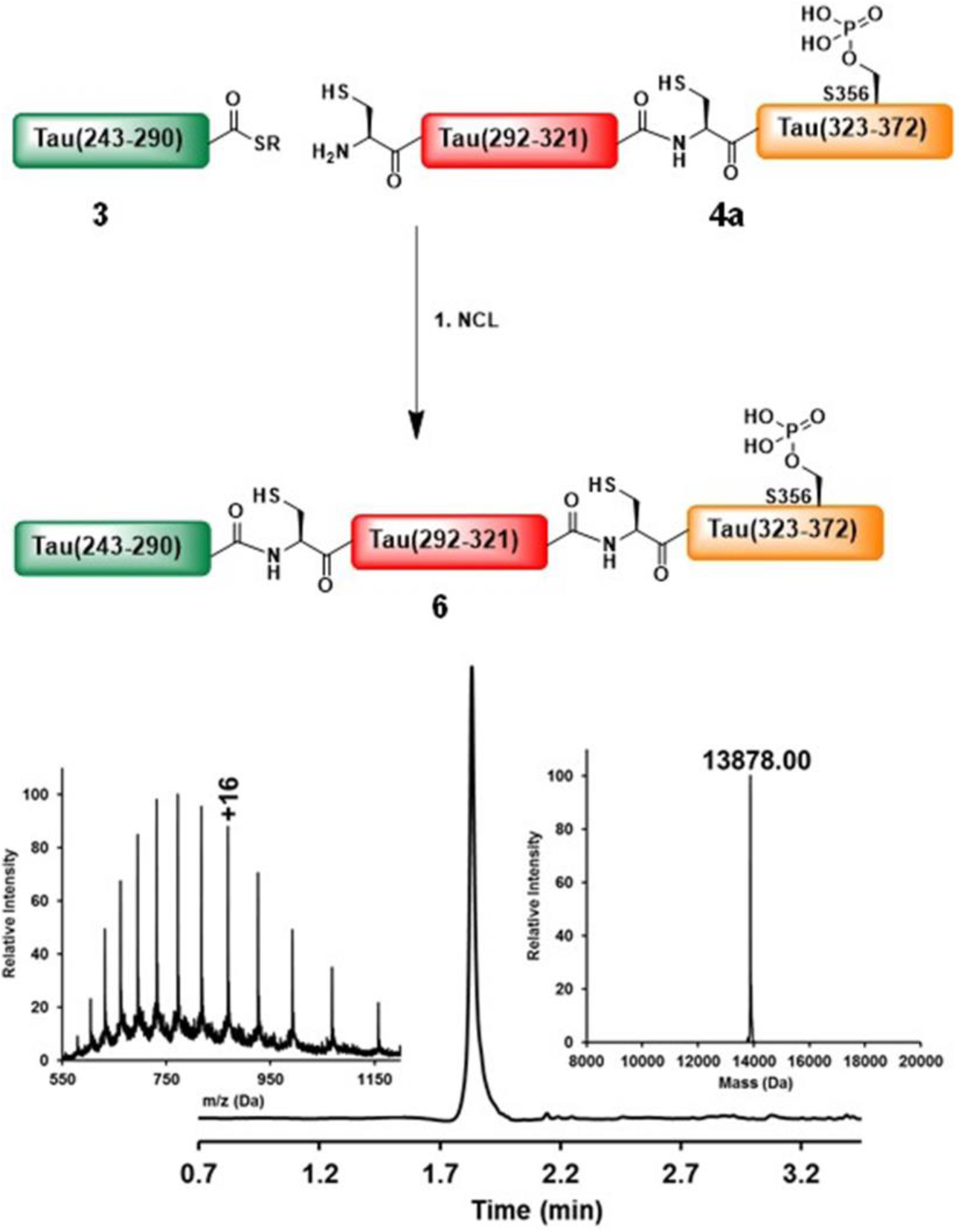
Native chemical ligation for the preparation of K18_pS356 (6). A) Schematic representation of the total synthesis of monophosphorylated tau (243-372, pS356) fragment (K18_pS356). B) Analytical RP-HPLC and ESI-MS spectra of K18_pS356, with the observed mass of 13878.00 Da (calculated 13875.89 Da).

### Preparation of diphosphorylated K18 at Ser 356 & Ser 262: Ligation of tau(243-290, pS262)-SR with tau(291-372, pS356)

**tau (291-372, pS356) (4a)** (2.3 mg, 1 eq) and **tau (243-290, pS262)-SR (3a)** (2.0 mg, 1.4 eq) were dissolved in NCL purged buffer (8.0 M urea, 50 mM TCEP, 2% TFET, pH 7.0). The mixture was incubated at 37 °C for 3 h with orbital agitation at 600 rpm and monitored by RP-UHPLC and mass spectrometry analyses. The ligated product was purified by RP-HPLC using a semipreparative C18 column with a linear gradient of 5-23% B (20 min), 20-40% B over 50 min, resulting in a 25.5% yield of product **7** (Fig. S11).

**Fig. S11.**
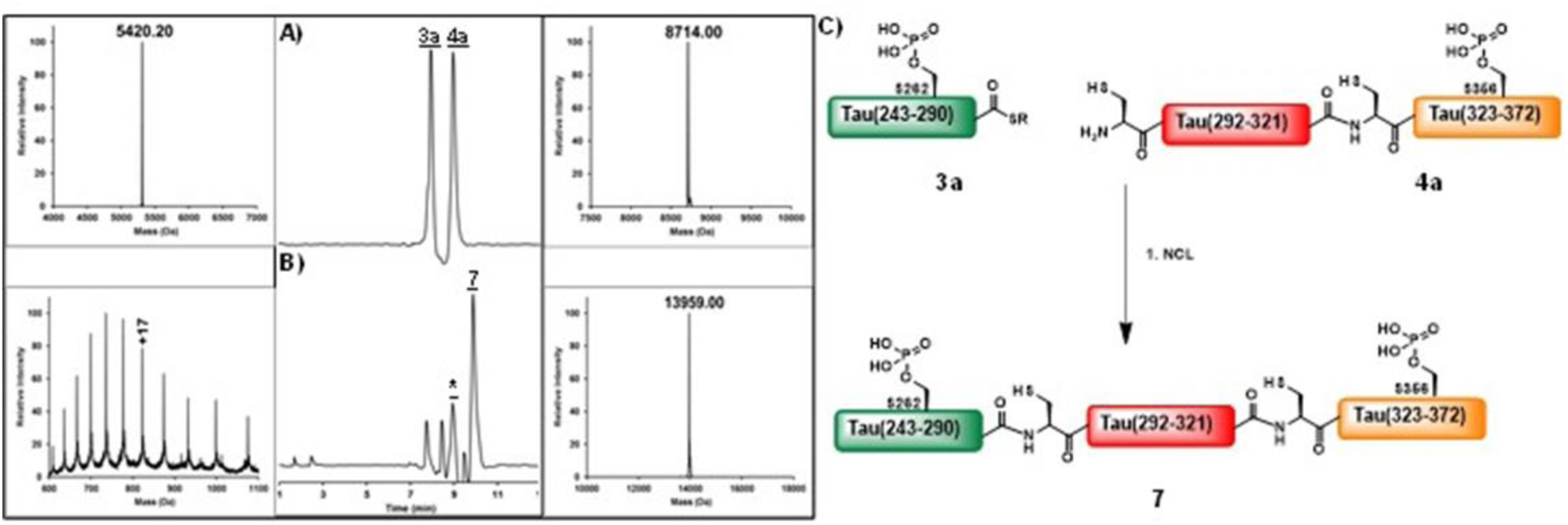
Native chemical ligation for the preparation of K18_pS356, pS262 (7). Analytical RP-HPLC and ESI-MS spectra of the ligation reaction between tau (291-372, pS356) and tau (243-290, pS262)-SR at A) 0 h and after B) 3 h. Peak **7** corresponds to the ligation product with the observed mass of 13959.00 Da (calculated 13955.85 Da). Peak * corresponds to the hydrolyzed thioester. C) Schematic representation of the total synthesis of diphosphorylated tau (243-372, pS356, pS262) fragment (K18_pS356, pS262).

### Preparation of triphosphorylated K18 at Ser 356, Ser 262, & Ser 258: Ligation of tau(243-290, pS262, pS258)-SR with tau(291-372, pS356)

**tau (291-372, pS356) (4’)** (2.8 mg, 1 eq) and **tau (243-290, pS262, pS258)-SR (3b)** (4.4 mg, 2.4 eq) were dissolved in NCL purged buffer (8.0 M urea, 50 mM TCEP, 2% TFET, pH 7.0). The mixture was incubated at 37 °C for 2 h with orbital agitation at 600 rpm and monitored by RP-UHPLC and mass spectrometry analyses. The ligated product was purified by RP-HPLC using a semipreparative C18 column with a linear gradient of 20-40% B over 40 min, resulting in a 27% yield of product **8** (Fig. S12).

**Fig. S12.**
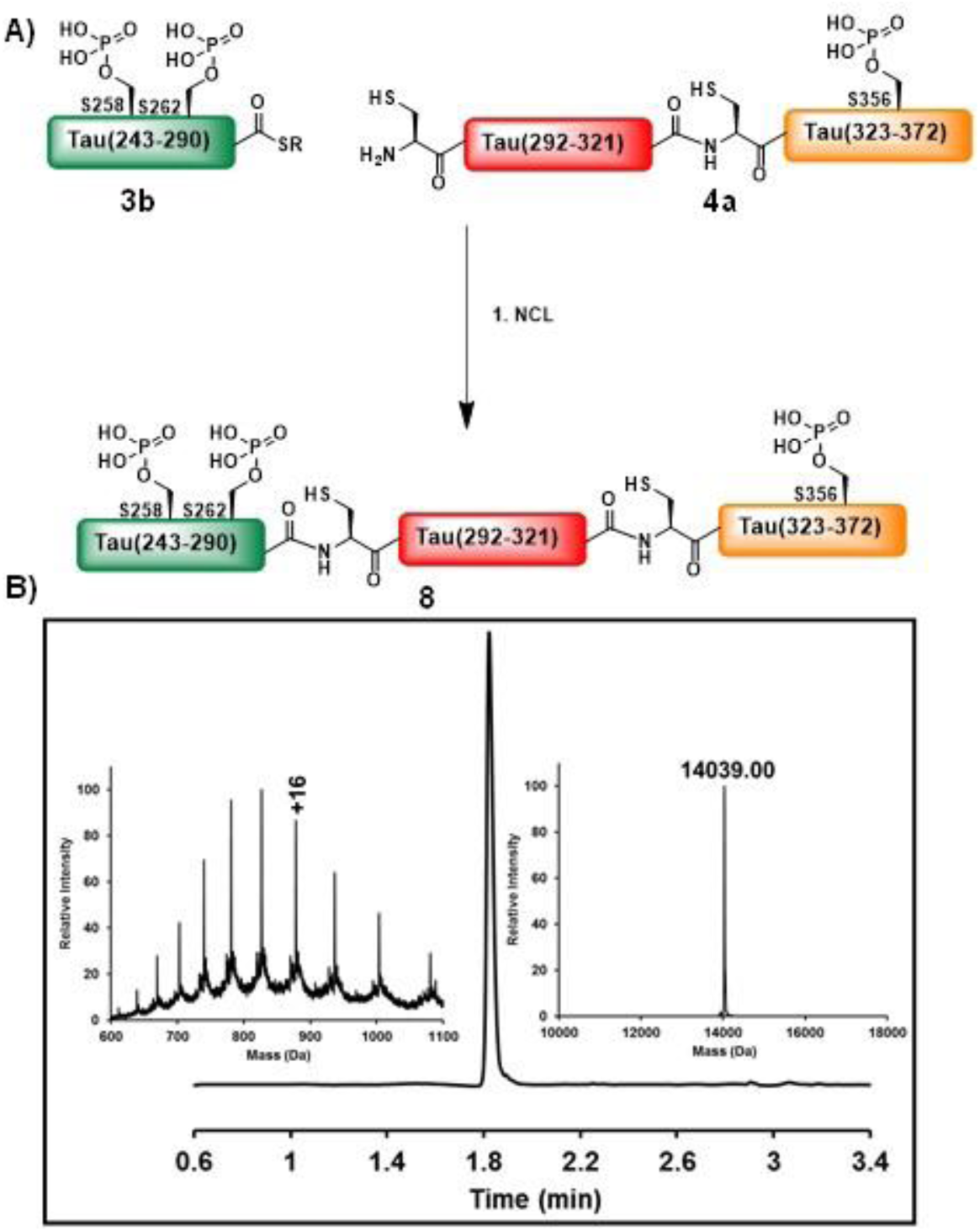
Native chemical ligation for the preparation of K18_pS356, pS262, pS258 (8). A) Schematic representation of the total synthesis of the triphosphorylated tau (243-372, pS356, pS262, pS258) fragment (K18_pS356, pS262, pS258). B) Analytical RP-HPLC and ESI-MS spectra of the purified K18 fragment triphosphorylated at Ser 356, Ser 262, and Ser 258, with the observed mass of 14039.00 Da (calculated 14035.89 Da).

### *In vitro* phosphorylation of tyrosine 310 (Tyr 310)

**K18_pS356, pS262, pS258** (1 mg, 1 eq) was dissolved in 10% urea and subsequently diluted in 50 mM Tris.HCl (pH 7.0) containing 1 mM DTT, 5 mM MgCl2, 20 mM Va3NO4, and 5 mM ATP for a final concentration of peptide of 1 mg/ml. The enzymatic assay was initiated by incubation of peptide **8** with c-Abl (0.7 mg, 0.2 eq) for 5 h at 30 °C without agitation. The reaction was monitored by RP-UHPLC and mass spectrometry analyses to verify the completion of phosphorylation. The phosphorylated peptide was purified by RP-HPLC using a semipreparative C18 column with a linear gradient of 20-35% B over 50 min, resulting in 50% yield of the tetraphosphorylated K18 peptide at residues Ser 356, Tyr 310, Ser 262 and Ser 258 (Fig. S13).

**Fig. S13.**
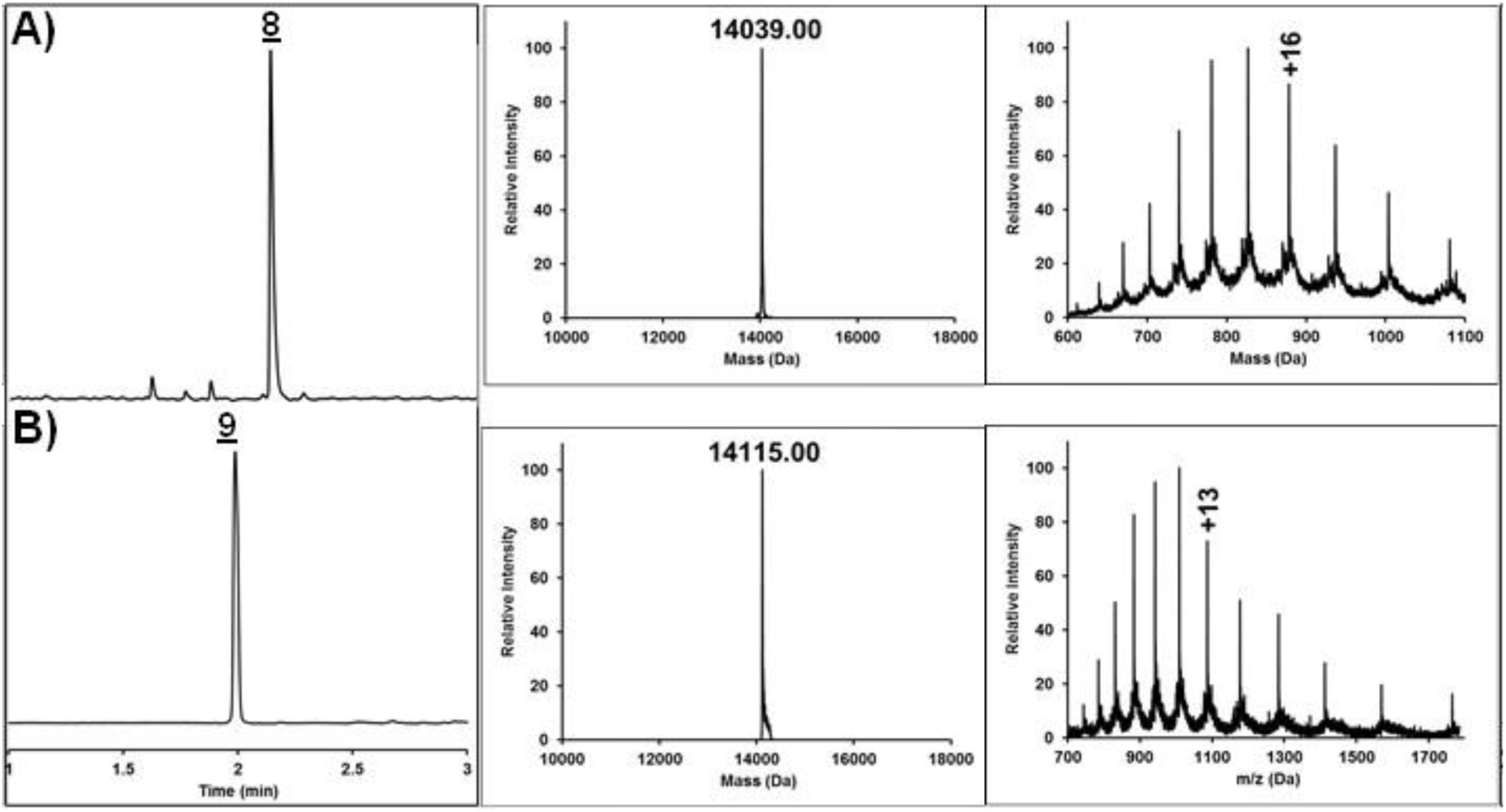
Analytical RP-HPLC and ESI-MS spectra of the *in vitro* phosphorylation reaction by c-Abl kinase at A) 0 h and after B) 5 h. Peak **9** corresponds to the tetraphosphorylated K18_pS356, pY310, pS262, pS258) with the observed mass of 14115.00 Da (calculated 14115.89 Da).

### Circular dichroism (CD) measurements

CD spectra were acquired on a Jasco J-815 CD spectrometer using a 1 mm quartz cuvette. Spectra were measured from 250 to 195 nm with a step size of 0.2 nm, a bandwidth of 1 nm, and a response time of 8 sec at 20 °C. For each spectrum, 5 scans were obtained and averaged. The averaged spectra were processed by smoothing using a binomial filter with a convolution width of 99 data points, and the resulting spectra were plotted as the mean residue molar ellipticity as a function of the wavelength.

### *In vitro* aggregation studies

Lyophilized protein samples were dissolved in aggregation buffer (10 mM phosphate, 50 mM NaF, pH 7.4) to a final concentration of 10 μM. Protein concentrations were determined using a NanoDrop 1000 spectrophotometer operated at 280 nm. The pH was adjusted to 7.4 using 0.1 M NaOH, followed by filtration through 100 kDa molecular weight cutoff (MWCO) filters. Ten microliters of each K18 construct was incubated at 37 °C in the presence of heparin sodium salt (Applichem GmbH, Germany) at a 1:4 molar ratio (heparin:protein) supplemented with 10 μM of ThS (T1892 Sigma). The extent of fibrillization was measured in triplicate on a FLUOstar Omega® plate reader as a function of the ThS fluorescence (excitation wavelength: 440 nm, emission wavelength: 480 nm, cycle time: 10 min).

### Transmission electron microscopy (TEM)

A total of 5 μL of each sample was deposited onto glow-discharged Formvar-coated 200 mesh copper grids (Electron Microscopy Sciences) for 90 sec. Grids were washed twice with dd H2O, stained with 0.7% (w/v) uranyl formate for 30 sec and dried. A Tecnai Spirit BioTWIN electron microscope operated at 80 kV and equipped with a LaB6 gun and a 4K × 4K FEI Eagle CCD camera was used to observe the samples and acquire the micrographs.

**Fig. S14.**
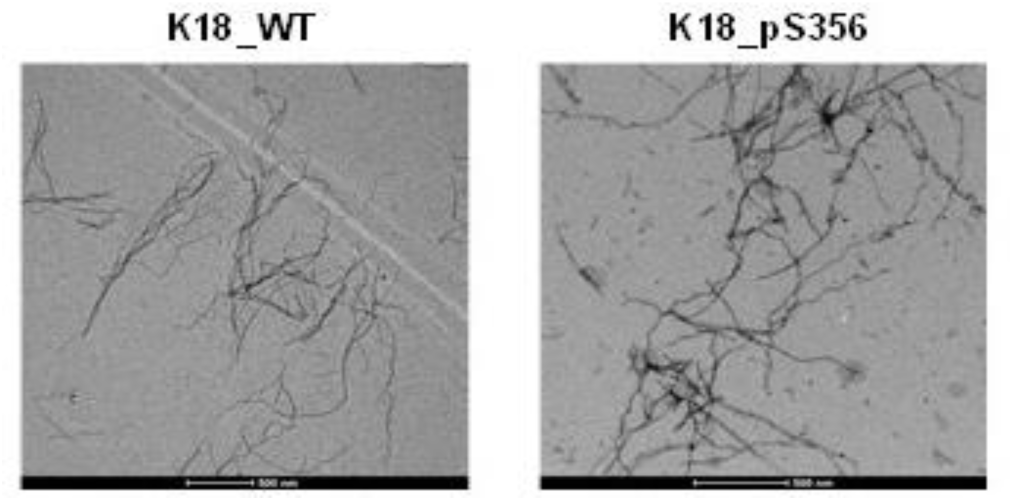
TEM images of WT and K18_pS356, after 48 h incubation at 37°C in the presence of heparin (scale bars are 500 nm).

### Seeding assays

#### Liposome-mediated transduction of tau seeds

HEK293T tau biosensor cells stably express the tau repeat domain containing the disease-associated mutation (P301S), fused to cyan fluorescent protein (CFP) or yellow fluorescent protein (YFP). Cells were plated at 35,000 cells per well in a 96-well plate. After 18 h, the cells reached 60% confluency and were transduced with WT, mono-, di- and triphosphorylated K18 aggregate seeds at a final concentration of 100 nM (monomer equivalent) per well. Transduction complexes were prepared by combining [8.75 μL Opti-MEM (Gibco) +1.25 μL Lipofectamine 2000 (Invitrogen)] with [Opti-MEM + aggregate seeds] for a total volume of 20 μL per well. Cells were incubated with the transduction complexes for 48 h to induce intracellular aggregation.

#### FRET flow cytometry

Cells were harvested with 0.05% trypsin and fixed in 2% paraformaldehyde (Electron Microscopy Services) for 10 min, after which they were resuspended in flow cytometry buffer. A MACSQuant VYB (Miltenyi) was used to perform FRET flow cytometry. To measure CFP and FRET, cells were excited with a 405 nm laser, and fluorescence was captured with 405/50 nm (CFP) and 525/50 nm filters (FRET). To measure YFP, cells were excited with a 488 nm laser, and fluorescence was captured with a 525/50 nm filter. To quantify FRET, we used a gating strategy similar to that previously described. The integrated FRET density (IFD), defined as the percent of FRET-positive cells multiplied by the median fluorescence intensity of the FRET-positive cells, was used for all analyses. For each experiment, 20,000 cells were analyzed in quadruplicate. Analysis was performed using FlowJo v10 software (Treestar).

#### Seeding experiment 1

K18 tau variants were incubated with or without heparin (Sigma) for a period of 24 h or 72 h at 25°C or 37°C. Samples were then used in the seeding assay at a final concentration of 100 nM (monomer equivalent) per well. Error bars represent the standard deviation.

**Fig. S15.**
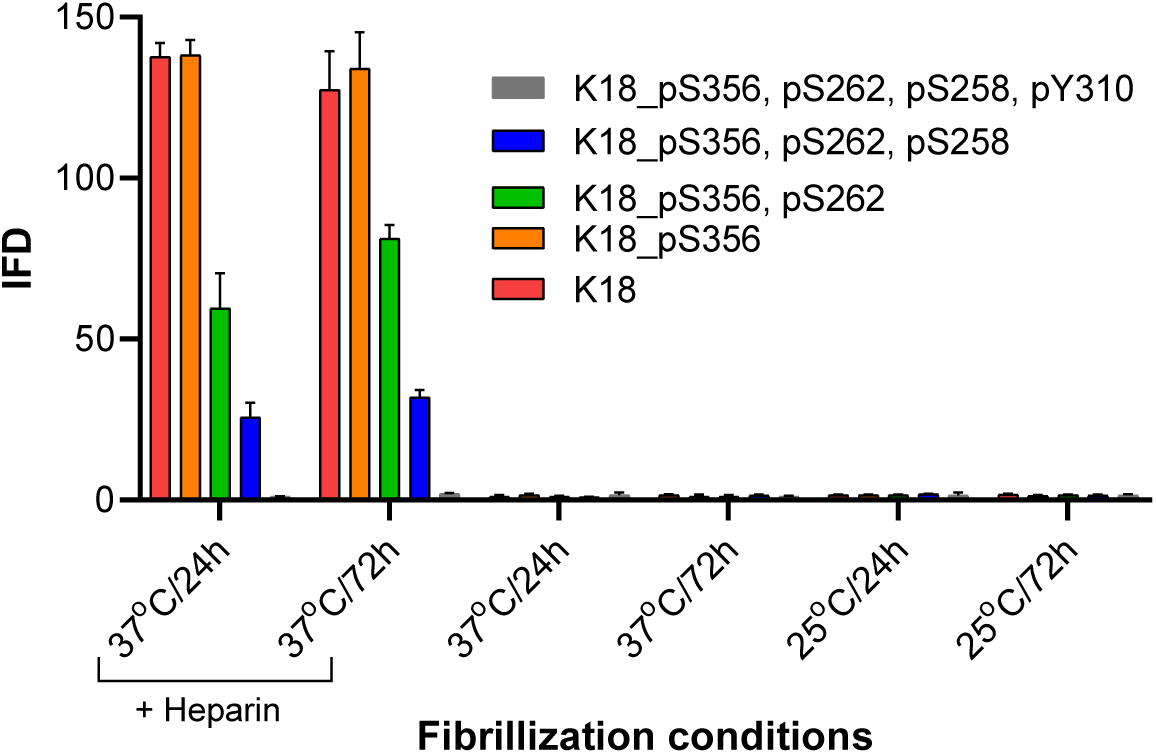
IFD measurements following the transduction of the WT, mono-, di- and triphosphorylated K18 previously incubated with or without heparin for various incubation times and temperatures.

#### Seeding experiment 2

All constructs, except for K18_pS356, pS262, pS258, pY310, were incubated with heparin for various time periods (0, 30 min, and 1, 2, 3, 6, 12, 24 h) at 37°C. The samples were then frozen in liquid nitrogen and kept at -80°C. To quantify the seeding activity, samples were thawed on ice and tested immediately at 100 nM (monomer equivalent) per well. Error bars represent the standard deviation.

#### Seeding experiment 3

K18 and K18_pS356 were incubated with heparin for various time periods (0, 1, 3, 5, 10, 15, 30, 45, 60 and 120 min) at 37°C. They were then frozen in liquid nitrogen and maintained at -80°C. To quantify the seeding activity, the samples were thawed on ice and tested immediately at 100 nM (monomer equivalent) per well. Error bars represent the standard deviation.

**Fig. S16.**
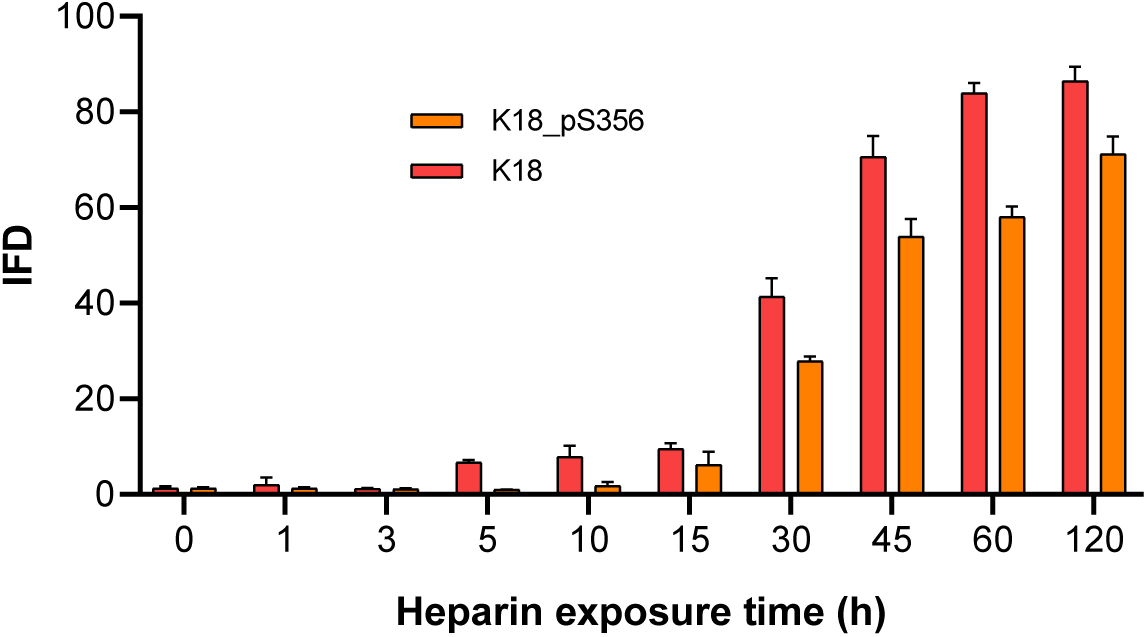
IFD measurements following the transduction of WT and pS356 K18 previously incubated at 37°C in the presence of heparin for various incubation times (0, 1, 3, 5, 10,15, 30, 45, 60 and 120 min).

#### Tubulin polymerization assay

The tubulin assembly assay for K18 or the phosphorylated K18 (K18_pS356, K18_pS262, pS356, or K18_pS262, pS258, pS356) proteins was initiated by mixing 30 μM protein with tubulin (40 μM, in 100 μl total volume) in MT assembly buffer (80 mM PIPES pH 6.9, 2 mM MgCl2 and 0.5 mM EGTA) supplemented with 1 mM GTP at 4 °C. The samples were transferred and incubated in a water bath set at 37 °C for 4 min. Then, the polymerization was monitored by measuring the absorbance at 350 nm every 30 sec using a FLUOstar Omega plate reader incubated at 37 °C. Each experiment was performed in triplicate and repeated at least two times.

